# Mechanism and effects of the skeletal muscle Nav1.4 inhibition by cannabidiol

**DOI:** 10.1101/2020.06.30.180943

**Authors:** Mohammad-Reza Ghovanloo, Koushik Choudhury, Tagore S. Bandaru, Mohamed A. Fouda, Kaveh Rayani, Radda Rusinova, Tejas Phaterpekar, Karen Nelkenbrecher, Abeline R. Watkins, Damon Poburko, Jenifer Thewalt, Olaf S. Andersen, Lucie Delemotte, Samuel J. Goodchild, Peter C. Ruben

**Author notes:** Corresponding Author: Dr. Peter C. Ruben, Department of Biomedical Physiology and Kinesiology, Simon Fraser University, 8888 University Drive, Burnaby, BC, Canada V5A 1S6. Indicates these authors contributed equally (Koushik Choudhury, Tagore S. Bandaru, Mohamed A. Fouda, Kaveh Rayani).

## Abstract

*Cannabis sativa* contains active constituents called phytocannabinoids. Some phytocannabinoids are psychotropic and others are not. The primary non-psychotropic phytocannabinoid is cannabidiol (CBD), which is proposed to be therapeutic against many conditions, including muscle spasms. Mechanisms have been proposed for the action of CBD on different systems, involving multiple targets, including the voltage-gated sodium channel (Nav) family, which are heavily implicated in many of the conditions CBD has been reported to relieve. In this study, we investigated the modulatory mechanism of CBD on Nav1.4. Based on previous results, we tested the hypothesis that CBD mechanism of action involves: 1) modulation of membrane elasticity, which indirectly contributes to Nav inhibition; and 2) physical block of the Nav pore. We first performed molecular dynamic (MD) simulations to visualize CBD effects and localization inside the membrane, and then performed NMR to verify the MD results, showing CBD localizes below membrane headgroups. Then, we performed a gramicidin-based fluorescence (GFA) assay that showed CBD alters membrane elasticity. Next, we used site-directed mutagenesis in (F1586A) and around (WWWW) the Nav1.4 pore. Removing the local anesthetic binding site with F1586A reduced CBD block of INa. Occluding the fenestrations with WWWW blocked CBD access from the membrane into the Nav1.4 pore. However, stabilization of inactivation, via CBD-induced changes in membrane elasticity persisted, in WWWW. To investigate the potential therapeutic value of CBD against some Nav1.4 channelopathies, we used a pathogenic variant of Nav1.4, P1158S, known to cause myotonia and periodic paralysis. We found CBD reduces excitability in both wild-type and the mixed myotonia/periodic paralysis variant. Our *in-vitro/in-silico* results suggest that CBD may have therapeutic value against myotonia. Because Nav1.4 is crucial to skeletal muscle contraction, we used rat diaphragm myography and found the presence of saturating levels of CBD reduces skeletal muscle contraction.

**SUMMARY:** We used multidisciplinary approaches to show the mechanism and pathway by which CBD inhibits the skeletal muscle, Nav1.4. Our results suggest CBD modulates membrane elasticity and directly interacts with Nav1.4 within its pore.

## INTRODUCTION

The cannabis plant, *Cannabis sativa*, contains over 120 active constituents, collectively known as phytocannabinoids(Morales et al., 2017). Some phytocannabinoids mediate psychotropic effects, whereas others do not(Morales et al., 2017). The primary non-psychotropic phytocannabinoid is cannabidiol (CBD). The structure of CBD is nearly identical to the main psychotropic compound isolated from cannabis, Δ9-tetrahydracannabinol (THC)(Morales et al., 2017). The only structural difference between the two compounds is the presence of a free hydroxyl in CBD in place of a closed ring in THC. This structural difference underlies THC’s high affinity for the human cannabinoid receptors, CB1 and CB2, thought to mediate the euphoria associated with using cannabis(Devinsky et al., 2017). In contrast to THC, CBD has little to no affinity for CB receptors(Pertwee, 2008). However, CBD has been suggested to be a potentially therapeutic compound against a variety of different conditions, including muscle spasms, pain, and seizures. Some reports of CBD efficacy are anecdotal, whereas others have been experimentally and clinically substantiated(Devinsky et al., 2017). CBD showed therapeutic efficacy in a recent phase III human clinical trial against Dravet syndrome(Devinsky et al., 2017), a severe form of childhood epilepsy and received FDA approval for its treatment.

Reports of CBD efficacy, along with its low affinity for CB receptors, have inspired studies on its CB-independent actions. Many mechanisms and targets have been proposed for the action of CBD(Kaplan et al., 2017; Ghovanloo et al., 2018c; Ross et al., 2008; De Petrocellis et al., 2011; Patel et al., 2016). The voltage-gated sodium channel (Nav) family is among these suggested targets(Patel et al., 2016; Ghovanloo et al., 2018c), in part because Nav underpin many of the conditions for which CBD has been shown, or suggested, to be efficacious.

Sodium currents through Nav initiates action potentials (AP) in neurons, myocardium, and skeletal muscles. Nav are hetero-multimeric proteins composed of a large ion conducting and voltage-sensing α-subunit and smaller β-subunits(Ghovanloo et al., 2016; Cannon, 2006; Ghovanloo and Ruben, 2020; Jiang et al., 2020; Catterall, 2014). The α-subunit is a single transcript that includes four 6-transmembrane segment domains. Each structural domain can be divided into two functional sub-domains: the voltage-sensing domain (VSD) and the pore-domain (PD)(Ghovanloo et al., 2016). These functional sub-domains are connected through the intracellular S4-S5 linker(Yarov-Yarovoy et al., 2012). The Nav pore is the site of interaction for many pharmacological blockers(Lee et al., 2012; Gamal El-Din et al., 2018). The pore is surrounded by four intra-bilayer fenestrations whose functional roles remain speculative(Pan et al., 2018).

Variants of the Nav subtype predominantly expressed in skeletal muscles, Nav1.4, are associated with contractility dysfunction. Most Nav1.4 variants depolarize the sarcolemma; this depolarization can result in either hyper- or hypo-excitability in phenotypes(Cannon, 2006). Hyperexcitable muscle channelopathies are classified as either non-dystrophic myotonias or periodic paralyses(Lehmann-Horn et al., 2008). Most of these channelopathies arise from sporadic *de-novo* or autosomal dominant mutations in *SCN4A*(Ghovanloo et al., 2018a).

The majority of gain-of-function (GOF) Nav1.4 variants result in myotonic syndromes, which are defined by a delayed relaxation after muscle contraction(Lehmann-Horn and Rudel, 1995; Tan et al., 2011). In myotonia, there is an increase in muscle membrane excitability in which even a brief voluntary contraction can lead to a series of APs that can persist for several seconds after motor neuron activity is terminated, a condition that is perceived as muscle stiffness(Tan et al., 2011). The global prevalence of non-dystrophic myotonias is ∼1/100,000(Emery, 1991). This condition is not considered lethal, but it can be life-limiting due to the multitude of problems it can cause, including stiffness and pain(Vicart et al., 2005).

A cationic leak (gating-pore current in the VSD) with characteristics similar to the ω-current in *Shaker* potassium channels has been shown to cause periodic paralyses(Jiang et al., 2018). This mechanism indicates that periodic paralyses can be caused by a severe form of GOF in Nav1.4(Wu et al., 2011; Tombola et al., 2005). A subset of periodic paralyses is triggered by low serum [K^+^] and results in episodes of extreme muscle weakness. These are known as hypokalemic periodic paralyses (hypoPP)(Miller et al., 2004). hypoPP onsets between the ages of 15– The prevalence of hypoPP is also ∼1/100,000. A serum [K^+^] less than 3 mM (normal concentration=3.5–5.0) may trigger hypoPP(Fontaine, 2008).

There are few therapeutics for these skeletal muscle dysfunctions, and treating myotonias and periodic paralyses mostly relies on drugs developed for other conditions, including local-anesthetics (LA). Myotonia treatment is focused on reducing the involuntary AP bursts(Vicart et al., 2005; Desaphy et al., 2004); hypoPP treatment is mostly focused on restoring serum [K^+^](Torres et al., 1981; Tawil et al., 2000; Sternberg et al., 2001; Venance et al., 2004). Thus, there remains a need for compounds that alleviate the hyperexcitability associated with both myotonia and hypoPP. Interestingly, non-euphoric plant cannabinoids have been shown to enhance muscle quality and performance of dystrophic mdx mice(Iannotti et al., 2019).

In this study, we first sought to delineate the mechanisms by which CBD may affect Nav1.4. We previously described the inhibitory effects of CBD on some neuronal Nav subtypes, which prompted us to predict a possible mechanism in which CBD both directly (direct interaction with channel) and indirectly (modulating membrane elasticity) inhibits Nav(Ghovanloo et al., 2018c). In the present study, we explored whether CBD accumulates in the membrane, which could alter membrane elasticity, and, while residing inside the membrane, enters the Nav1.4 fenestrations and blocks the channel pore(Gamal El-Din et al., 2018). Next, we explored the effects of CBD on a mixed periodic paralysis and myotonia Nav1.4 mutation (P1158S)(Ghovanloo et al., 2018a; Webb and Cannon, 2008) to determine whether it could alleviate the mutant phenotypes. Finally, we sought to survey whether saturating levels of CBD can reduce skeletal muscle contractility, which it does.

## RESULTS

### Molecular dynamics (MD) simulations predict CBD accumulates in the hydrophobic region of phospholipid bilayers

We previously determined that CBD non-selectively inhibits voltage-dependent sodium and potassium currents with a steep average Hill-slope of ∼3, which suggested multiple interactions. Contrary to what is expected for classic pseudo–second order bimolecular blocking schemes, we found CBD was fastest to equilibrate and most potent at lower temperatures(Ghovanloo et al., 2018c). These findings, together with CBD’s stabilizing effects on neuronal Nav inactivation and CBD’s high LogP of ∼5.9, led us to explore whether CBD alters membrane elasticity, which indirectly could inhibit Nav currents(Ghovanloo et al., 2018c), similar to what has been suggested for amphiphilic compounds(Lundbæk et al., 2004; Lundbaek, 2005). To test this hypothesis, we performed MD simulations of CBD (in mM concentration in membrane) on 1-palmitoyl-2-oleoyl-sn-glycero-3-phosphocholine (POPC) lipid membranes in the hundreds of nanoseconds range (**Figure 1a-e; Table 1**). MD results indicate that in both symmetrical (i.e. the same number of CBD molecules in both leaflets of the membrane) and asymmetrical (i.e. CBD in a single leaflet), there are no substantial changes in the area per lipid.

**Table 1.** Details of the simulation systems.

| **System** | **Number of**<br>**CBD**<br>**molecules** | **Number of**<br>**lipid**<br>**molecules** | **Number of**<br>**water**<br>**molecules** | **Number of**<br>**ions (Na/Cl)** | **Total**<br>**number of**<br>**atoms** |
| --- | --- | --- | --- | --- | --- |
| #0 | 0 | 188 | 11377 | 30/30 | 59383 |
| #1 | 2 (symmetric) | 178 | 11348 | 30/30 | 58062 |
| #2 | 3 (asymmetric) | 171 | 11377 | 30/30 | 57264 |
| #3 | 6 (symmetric) | 171 | 11377 | 30/30 | 57423 |

**Figure 1.**
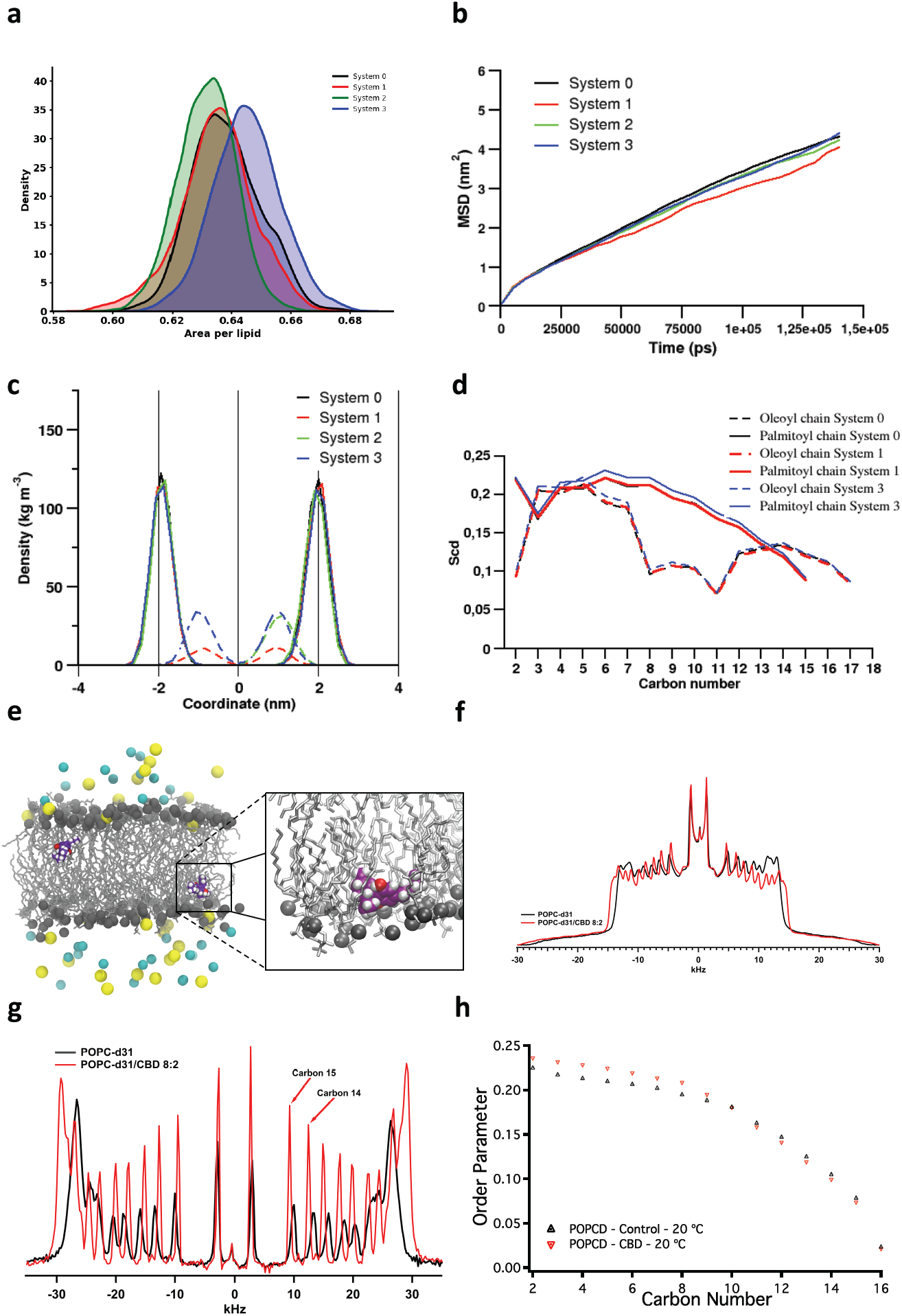
Effects of CBD on POPC membrane, via MD simulations and H^2^ NMR. (**a-b**) The effects of CBD on POPC membrane area per lipid and lipid diffusion. System 0 is control, System 1 is 2 CBD molecules in symmetry (1 in each leaflet), System 2 is 3 CBD molecules in asymmetry (only in a single leaflet), System 3 is 6 CBD molecules in symmetry (3 in each leaflet). (**a**) Area per lipid and (**b**) mean square displacement as a function of time are not affected by CBD. (**c**) Distribution of CBD into the membrane across a range of conditions. The distribution of phosphate groups is shown as solid lines, the distribution of CBD as dotted lines. The bilayer thickness remains ∼4 nm in presence and absence of CBD. (**d**) Order parameter of lipid acyl chains estimated from the MD simulations. (**e**) Snapshot of a CBD molecule in the POPC leaflet extracted from the MD simulations (see **Movie S1**). The zoomed-in image shows localization of CBD molecule below the leaflet headgroup. (**f**) NMR spectra collected at 20°C. (**g**) De-packed traces. (**h**) Order parameter calculation from NMR.

**Figure 1c** shows CBD density estimates as a function of membrane leaflet coordinate, where the lipid bilayer is centered at 0 (membrane thickness of ∼4 nm is unchanged across conditions). These results show that, in symmetrical conditions, there are two density peaks in both negative and positive coordinate ranges with an almost perfect overlap, and that CBD localizes in the area between the lipid headgroups and the membrane center, close to the lipid headgroup region. In the asymmetric condition, with 3 CBD molecules initially placed in the leaflet to the right, there is only a single peak in the positive coordinate range. The MD results show that CBD molecules tend to reside in the leaflet to which they originally were added, where they interact with, and detach rapidly from polar residues at the bilayer/solution interface and occasionally, move toward the water molecules outside the lipid. However, CBD then quickly moves back down into the lipid (**Movie S1**). This suggests, within the hundreds of nanoseconds timeframe of our simulations, that CBD does not diffuse across the two leaflets but, instead, tends to localize in the leaflet where it was initially placed. This could be a product of the oxygen atoms in CBD keeping it from diffusing across leaflets, and the CBD’s hydrophobic tail keeping it from getting too close to water molecules outside the membrane.

### ^2^H NMR verifies the MD predictions regarding localization

Next, we performed acyl chain order parameter calculations (from MD) which suggested that CBD causes a slight ordering of the membrane methylenes in the plateau region of the palmitoyl chain (C3-C8) (**Figure 1d**). Overall, the MD results suggest that CBD preferentially localizes under the phosphate heads, close to carbons 3-7 of the aliphatic chains of the POPC molecules (**Figure 1e**).

We tested MD predictions regarding CBD localization using NMR(Lafleur et al., 1989) with POPC-d31 and POPC-d31/CBD at a 4:1 ratio in deuterium depleted water at three different temperatures (20, 30, and 40°C) (**Figure 1f-h; Figure S1**). The NMR results were in striking agreement with the MD predictions of the changes in acyl chain order parameters and suggested that CBD causes an ordering of the C2-C8 methylenes, and a slight disordering from C10-C15, in a temperature-dependent manner.

### CBD alters bilayer elasticity in gramicidin-based fluorescence assay (GFA)

To explore CBD’s possible effects on lipid bilayer properties (membrane elasticity) that were predicted from the MD and NMR experiments, we tested CBD’s effects on lipid bilayer properties at concentrations where CBD has acute effects on Nav channels using a gramicidin-based fluorescence assay (GFA) which takes advantage of the gramicidin channels’ unique sensitivity to changes in bilayer properties(Andersen and Koeppe, 2007). The GFA is based on the gramicidin channels’ permeability to Tl^+^, a quencher of the water-soluble fluorophore 8-aminonaphthalene-1,3,6-trisulfonate (ANTS), which can be encapsulated in large unilamellar vesicles (LUVs) that have been doped with gramicidin. The rate of Tl^+^ influx, the rate of fluorescence quench, is a measure of the time-averaged number of gramicidin channels in the LUV membrane(Ingólfsson et al., 2010). Molecules that alter the thickness and elasticity of the LUV membrane will alter the lipid bilayer contribution to the free energy of dimerization, and thus the free energy of dimerization:

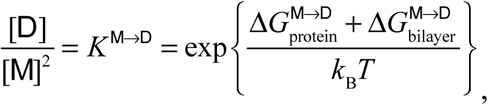

which will produce a shift in the gramicidin monomer*↔*dimer equilibrium (**Figure 2a**). Changes in this equilibrium will result in changes in the rate of Tl^+^ influx (fluorescence quench). CBD reduced the Tl^+^ influx rates in concentration-dependent manner (**Figure 2**). For comparison, we also show results obtained with Triton X-100, from(Ingólfsson et al., 2010), which increased the quench rates demonstrating that these two molecules have opposite effects on the membrane, and we conclude that CBD increases bilayer stiffness or thickness, whereas Triton X-100 decreases bilayer stiffness or thickness. Given that CBD has minimal effects in bilayer thickness (**Figure 1**), we conclude that CBD indeed alters lipid bilayer elasticity.

**Figure 2.**
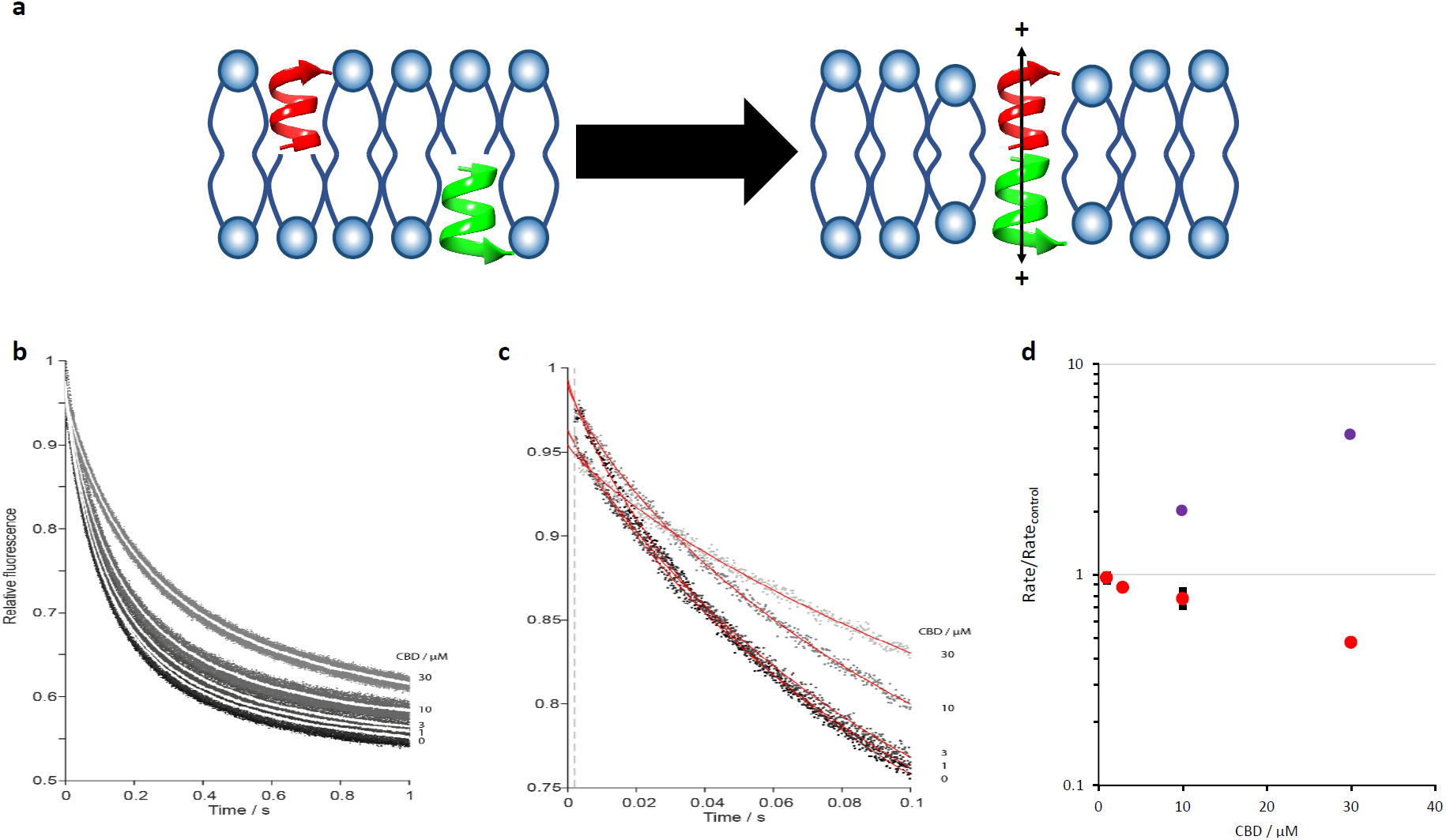
CBD alters lipid bilayer properties in gramicidin-based fluorescence assay (GFA). (**a**) Cartoon representation of gramicidin monomers in each leaflet coming together (dimerizing) to form cationic channels. The dimerization of the gramicidin channels is directly related to membrane elasticity. These properties are used to assay compound (e.g. CBD) effects on membrane elasticity. (**b**) Fluorescence quench traces showing Tl^+^ quench of ANTS fluorescence in gramicidin-containing DC_22:1_PC LUVs with no drug (control, black) and incubated with CBD for 10 min at the noted concentrations. The results for each drug represent 5 to 8 repeats (dots) and their averages (solid white lines). (**c**) Single repeats (dots) with stretched exponential fits (red solid lines). (**d**) Fluorescence quench rates determined from the stretched exponential fits at varying concentrations of CBD (red) and Triton X-100 (purple, from (Ingólfsson et al., 2010)) normalized to quench rates in the absence of drug. Mean ± SD, n = 2 (for CBD).

### CBD interacts with the Nav local-anesthetic site

We previously found that CBD displays an approximately 10-fold state-dependence (10-fold increased affinity for inactivated state) in Nav inhibition, a property similar to classic pore-blockers(Ghovanloo et al., 2018c; Kuo and Bean, 1994; Bean et al., 1983), which has also been observed with bilayer-modifying molecules(Lundbaek, 2005). Therefore, in that study, we tested CBD inhibition from the inactivated-state in a Nav1.1 pore-mutant (F1763A–LA mutant)(Ghovanloo et al., 2018c) and the results suggested a relatively small ∼2.5-fold drop in potency(Ghovanloo et al., 2018c). Here, to further explore possible CBD interactions at the pore, we performed molecular docking using the human Nav1.4 cryo-EM structure(Pan et al., 2018). **Figure 3a-b** show CBD docked onto the Nav1.4 pore, supporting a possible interaction at the LA site.

**Figure 3.**
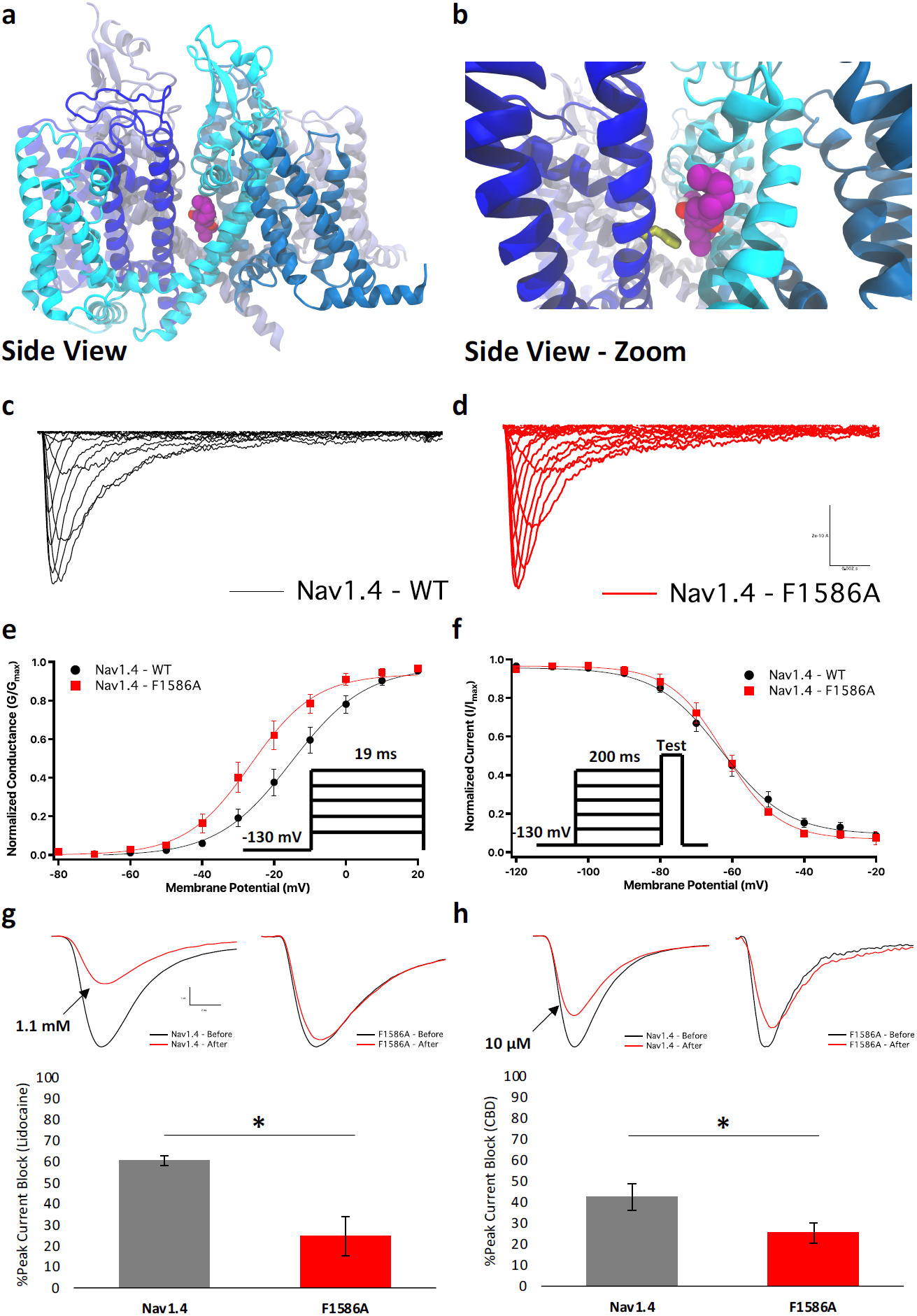
Inhibition of Nav1.4 pore by CBD, F1586A reduces inhibition. (**a**)Side-view of CBD docked into the pore of the human Nav1.4 structure. The structure is coloured by domain. DIV is coloured in deep blue. (**b**) Zoomed-in side-view, F1586 is coloured in yellow. (**c-d**) Representative families of macroscopic current traces from WT-Nav1.4 and F1586A. (**E**) Voltage-dependence of activation as normalized conductance plotted against membrane potential (Nav1.4: V_1/2_ = -19.9 ± 2.7 mV, z = 2.8 ± 0.3; F1586A: V_1/2_ = -22.4 ± 2.2 mV, z = 3.0 ± 0.3; n = 5-7). (**f**) Voltage-dependence of SSFI as normalized current plotted against membrane potential (Nav1.4: V_1/2_ = -66.9 ± 2.8 mV, z = -2.6 ± 0.3; F1586A: V_1/2_ = -63.3 ± 3.0 mV, z = -3.5 ± 0.3; n = 8-9). (**g-h**) Lidocaine/CBD inhibition of Nav1.4 and F1586A from -110 mV (rest) at 1 Hz (Lidocaine-Nav1.4: Mean block = 60.6 ± 2.3%; Lidocaine-F1586A: Mean block = 24.6 ± 9.3%; CBD-Nav1.4: Mean block = 42.4 ± 6.4%; CBD-F1586A: Mean block = 25.3 ± 4.8%; n = 3-5). Sample traces before and after compound perfusion are shown.

To test the docking result, we mutated the LA Nav1.4 F1586 into alanine (A) and performed voltage-clamp. **Figure 3c-f** show biophysical characterization of F1586A compared with WT-Nav1.4. We found that both channels have similar biophysical properties and, most importantly, the inactivation voltage-dependences were almost identical (p>0.05) (**Figure 3f**), suggesting that at any given potential both F1586A and Nav1.4 would have the same availability; therefore, pharmacological experiments could be performed using the same voltage-protocols on both channels.

In contrast to neuronal Navs that have inactivation midpoints (V1/2) of ∼-65 mV in neurons with resting membrane potentials (RMP) that are also ∼-65 mV, Nav1.4 has a V1/2 of ∼-67 mV in skeletal muscle fibers with an RMP of ∼-90 mV. This indicates that, whereas neuronal Navs are half-inactivated at RMP, Nav1.4 is almost fully available at RMP. Therefore, we measured lidocaine (positive control) and CBD inhibition of Nav1.4 from rest (−110 mV holding-potential to 0 mV, test-pulse at 1 Hz) to be closer to physiological conditions (**Figure 3g-h**). Our results suggest that 1.1 mM (resting IC50 on Nav1.4(Nuss et al., 1995)) lidocaine blocks ∼60% of INa in WT, and ∼20% in F1586A (p=0.020). 10 µM CBD blocks ∼45% INa in WT and ∼25% in F1586A (p=0.037). Hence, there is a 3-fold difference between lidocaine’s inhibition of WT vs. F1586A, and a smaller 1.5-fold difference for CBD inhibition. This suggests that while CBD may interact with the Nav pore similar to lidocaine, CBD’s interaction with F1586 is likely not as critical a determinant of its INa inhibition compared with lidocaine.

### CBD interacts with DIV-S6

Because CBD’s INa inhibition was less affected by F1586A, a mutation that destabilizes the LA binding-site, than a well-established pore-blocker like lidocaine, we investigated whether CBD interacts with the DIV-S6 (which includes F1586) or if it is inert, using isothermal titration calorimetry. We then compared CBD interactions to lidocaine. We found that both lidocaine and CBD appear to interact with the protein segment, though the nature of this interaction seems to differ (**Figure S2**). **Figures S2a-b** show sample ITC heat traces. Our results suggested that in the presence of protein, lidocaine titration causes an endothermic interaction. However, when the protein is absent, lidocaine titration into blank buffer causes exothermicity. In contrast, CBD titration in both blank buffer and in the presence of protein resulted in endothermicity. Interestingly, the magnitude of CBD’s heats of interaction were ∼4-fold larger in the protein condition compared to the blank condition. This was in contrast to lidocaine that showed comparable heats of interaction magnitudes in both conditions, but in different directions. To quantify interactions of both lidocaine and CBD, we subtracted the heats from runs with both protein and ligand subtracted from only ligand (blank). The subtracted heats show a similar trend between lidocaine and CBD (**Figure S2c-d**). These results suggest that both lidocaine and CBD interact with the protein segment; however, the nature of this interaction is different possibly due to a variation in physicochemical properties.

### CBD may penetrate into the pore through fenestrations

LAs block bacterial Navs in their resting-state by entering the pore through fenestrations in a size-dependent manner (i.e. smaller LAs get through more readily)(Gamal El-Din et al., 2018). Here, we sought to determine whether it is possible to block CBD’s access to the human Nav1.4 pore from the lipid phase of the membrane by occluding fenestrations. We previously found that CBD is highly lipid-bound (99.6%)(Ghovanloo et al., 2018c), and our MD results show that it preferentially localized in the hydrophobic part of the membrane, just below the lipid headgroups. Therefore, we reasoned that once CBD partitions into the membrane, it may access the Nav pore through the intramembrane fenestrations. To test this idea, we scrutinized the docking pose of CBD in the human Nav1.4 and observed its localization close to the fenestrations (**Figure 4a; Figure S3a-d**).

**Figure 4.**
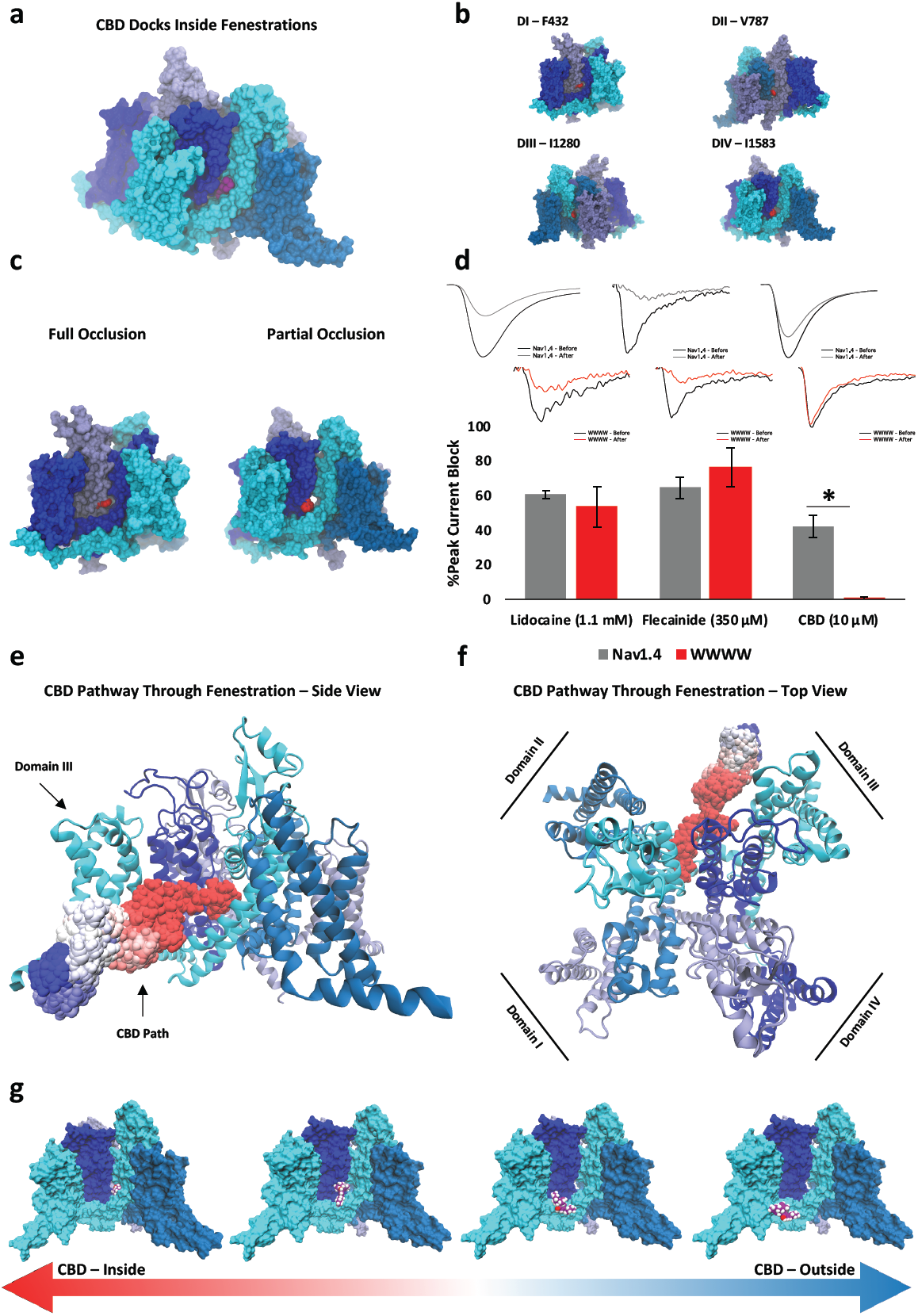
CBD interactions with and through Nav fenestrations. Side-view of CBD docked into the human Nav1.4 structure. The structure is coloured by domain (matched color to domain is shown in panel (**f**)), CBD is represented in purple. (**b**) Side-view of all four sides of human Nav1.4 (coloured by domain). Nav1.4 fenestrations are highlighted in red, along with the position of respective residues that were mutated into tryptophans (W). (**c**) Computational mutagenesis of fenestrations results 2 full and 2 partial occlusions (paralleled domains). (**d**) Lidocaine (1.1 mM) inhibition of Nav1.4 and WWWW from -110 mV (rest) at 1 Hz (Nav1.4: Mean block = 60.6 ± 2.3%; WWWW: Mean block = 53.6 ± 11.7%), flecainide (350 µM) inhibition (Nav1.4: Mean block = 64.6 ± 6.0%; WWWW: Mean block = 76.4 ± 11.3%), and CBD (10 µM) inhibition (Nav1.4: Mean block = 42.4 ± 6.4%; WWWW: Mean block = 6.4 ± 1.3%; n = 3-5 panel-wide). Traces before and after compound perfusion are shown. (**e**) CBD pathway through the Nav1.4 fenestration from side view, as predicted by MD simulations, red and blue correlate to CBD being inside and outside the fenestration, respectively (see **Movie S2-3**). (**f**) CBD pathway from top view of the channel. (**g**) Progressive snapshots of the movement of CBD over time from inside to outside the channel.

Next, we identified 4 residues (DI-F432, DII-V787, DIII-I1280, and DIV-I1583) that partially or fully occluded the fenestrations when mutated to tryptophan (W), as predicted by computational mutagenesis and structural minimization (partial versus full occlusion is due to structural asymmetry of mammalian Navs) (**Figure 4b-c**).

We measured resting-state block of 1.1 mM lidocaine, 350 µM flecainide, and 10 µM CBD from -110 mV on our WWWW construct. Our results suggest that lidocaine (p>0.05) and flecainide (p>0.05), but not CBD (p<0.05) blocked the WWWW mutant the same as WT (after compound has reached equilibrium) (**Figure 4d**). This is an interesting result considering that CBD is larger than lidocaine, but slightly smaller than flecainide. CBD was inert with respect to block of WWWW, relative to WT-Nav1.4, which suggests that CBD interacts with Nav1.4 via the fenestrations.

To visualize the possible pathway CBD follows through Nav1.4 fenestrations and into the pore at an atomistic resolution, we performed MD simulations in which we encouraged CBD to detach from its binding-site (see **Methods, Figure 4e-g; Figure S3e; Movie S2-3**). These results suggest that CBD approaches its binding-site in the pore via the fenestration without major reorganization of the channel structure.

### CBD does not affect Nav1.4 activation but stabilizes the inactivated state

We previously characterized the effects of CBD on Nav1.1 gating(Ghovanloo et al., 2018c). We found that CBD at ∼IC50 reduced channel conductance, did not change the voltage-dependence of activation, but produced a hyperpolarizing shift in steady-state fast inactivation (SSFI), and slowed recovery from fast (300 ms) and slow (10 s) inactivation(Ghovanloo et al., 2018c). Together with CBD’s inhibition of resurgent sodium currents(Patel et al., 2016; Ghovanloo et al., 2018c), these results suggested that CBD prevents the opening of Navs, but the channels that can open, activate with unchanged voltage-dependence and are more likely to inactivate. The overall effect is a reduction in excitability(Ghovanloo et al., 2018c). Here, we tested the hypothesis that CBD’s non-selectivity in INa inhibition suggests similar non-selectivity in modulating Nav gating (i.e. CBD imparts similar gating modulation across Nav subtypes). To test this idea, we assessed Nav1.4 activation in presence and absence of 1 µM CBD by measuring peak channel conductance at membrane potentials between −100 and +80 mV (**Figure 5a**). CBD did not significantly alter V1/2 or apparent valence (z) of activation (p>0.05). Normalized Nav1.4 currents as function of membrane potential are shown in **Figure 5b**. These results indicate that, as with Nav1.1, CBD does not alter Nav1.4 activation.

**Figure 5.**
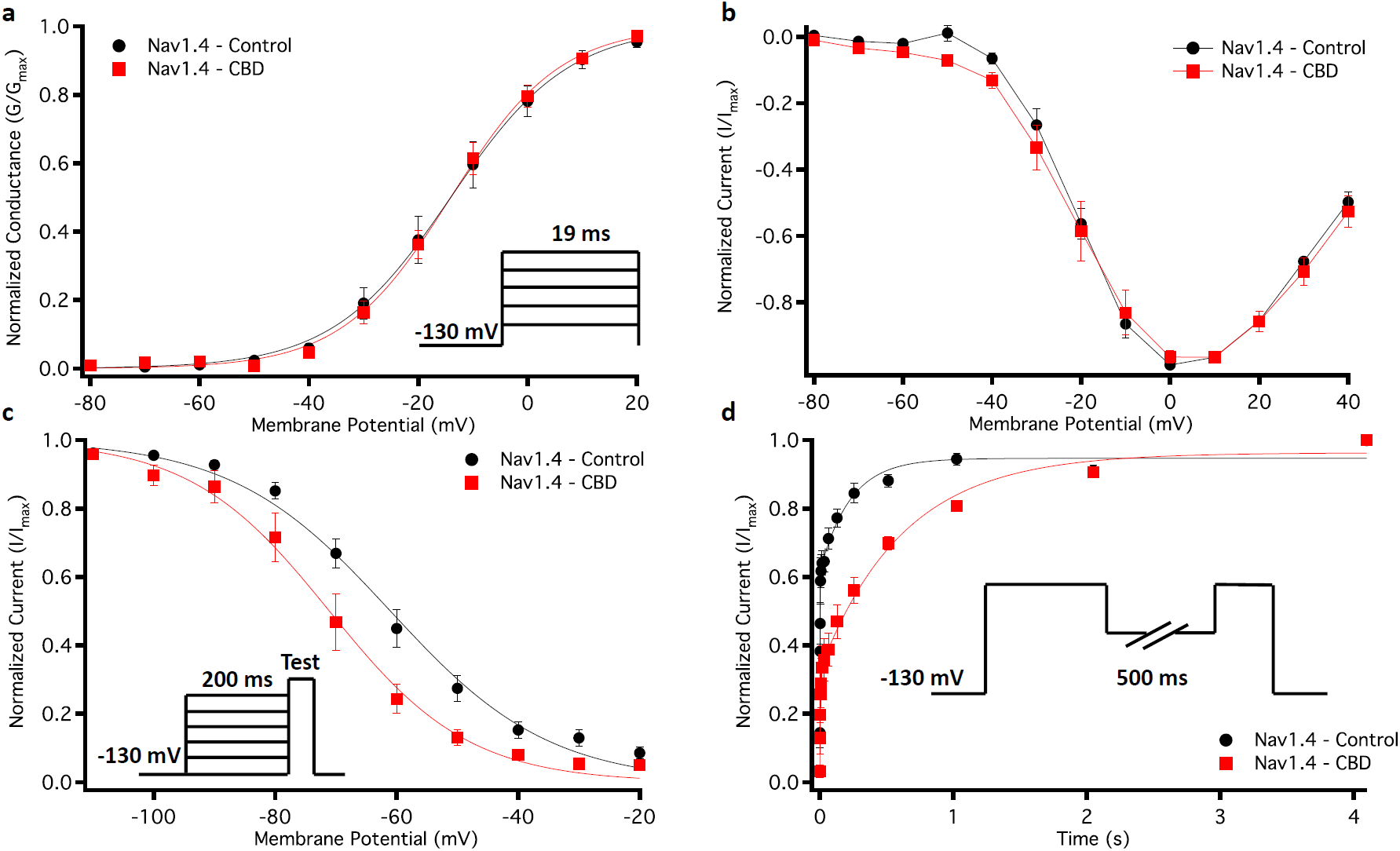
Effects of CBD (1 µM) on Nav1.4 gating. (**a-b**) Voltage-dependence of activation as normalized conductance plotted against membrane potential (Control: V_1/2_ = -19.9 ± 4.2 mV, z = 2.8 ± 0.3; CBD: V_1/2_ = -14.3 ± 4.2 mV, z = 2.8 ± 0.3; n = 5) and normalized activating currents as a function of potential. (**c**) Voltage-dependence of SSFI plotted against membrane potential (Control: V_1/2_ = -64.1 ± 2.4 mV, z = -2.7 ± 0.3; CBD: V_1/2_ = -72.7 ± 3.0 mV, z = -2.8 ± 0.4; n = 5-8). (**d**) Recovery from fast inactivation at: 500 ms (Control: *τ*_Fast_ = 0.0025 ± 0.00069 s, *τ*_Slow_ = 0.224 ± 0.046 s; CBD: *τ*_Fast_ = 0.0048 ± 0.00081 s; *τ*_Slow_ = 0.677 ± 0.054 s; n = 5-7).

Next, we examined the voltage-dependence of SSFI using a standard 200 ms pre-pulse voltage protocol. Normalized current amplitudes were plotted as a function of pre-pulse voltage (**Figure 5c**). These results mimicked our previous observations in Nav1.1(Ghovanloo et al., 2018c), in that CBD left-shifted the Nav1.4 inactivation curve (p<0.05).

To measure recovery from inactivation, we held Nav1.4 at -130 mV to ensure that the channels were fully available, then pulsed the channels to 0 mV for 500 ms and allowed different time intervals at -130 mV to measure recovery as a function of time. As previously observed in Nav1.1, CBD slowed the Nav1.4 recovery from inactivation (p<0.05), suggesting that it takes longer for CBD to come off the channels than the time it takes the channels to recover from inactivation (**Figure 5d**). Collectively, these results support our hypothesis that CBD non-selectively modulates Nav gating, and further suggests how CBD may reduce Nav1.4 excitability.

### CBD hyperpolarizes SSFI in Nav1.4-WWWW

To determine a possible association between membrane elasticity and stabilized inactivation, we measured effects of CBD before and after compound perfusion in the WWWW mutant, in a matched-pair manner. Although CBD did not inhibit peak INa, it hyperpolarized the SSFI curve (p<0.05), suggesting CBD’s modulation of membrane elasticity is at least in part responsible for stabilizing Nav inactivation (**Figure S4**). This is an interesting finding because our GFA results suggest that CBD increases bilayer stiffness or thickness, and previous studies suggest that compounds such as Triton X-100 that reduce this stiffness or thickness also hyperpolarize the Nav SSFI curve(Lundbæk et al., 2004).

### CBD affects a pH-sensitive mixed myotonia/hypoPP Nav1.4-mutant, P1158S (DIII-S4-S5)

Because CBD is therapeutic against seizure disorders(Devinsky et al., 2017), typically considered neuronal GOF conditions, we examined whether CBD may similarly ameliorate a skeletal muscle GOF condition(Cannon, 2015). We recently discovered that the P1158S mutation in Nav1.4 increases the channel’s pH-sensitivity(Ghovanloo et al., 2018a). The P1158S gating displays pH-dependent shifts that, using AP modeling, are predicted to correlate with the phenotypes associated with this variant. Therefore, the relationship between pH and P1158S could be used as an *in-vitro/in-silico* assay of Nav1.4 hyperexcitability (to model moderate to severe GOF). Here, we used this assay to investigate CBD’s effects on skeletal muscle hyperexcitability. We tested effects of 1 µM CBD (pK_a_=9.64) on P1158S at pH6.4 (myotonia-triggering) and pH7.4 (hypoPP-triggering). **Figure 6** shows CBD effects on P1158S at low and high pH. Interestingly, the lack of selectivity in gating modulation by CBD also exists in P1158S at both pHs. CBD did not change activation (p>0.05), but hyperpolarized inactivation (p<0.05) and slowed recovery from inactivation (p<0.05) (**Figure 6a-f**). Consistent with previous results where CBD inhibited persistent I_Na_(Ghovanloo et al., 2018c; Patel et al., 2016), CBD also reduced the exacerbated persistent I_Na_ associated with P1158S at pH7.4 (p<0.05) (**Figure 6g**). Persistent I_Na_ reduction could not be detected at pH6.4 (p>0.05) (**Figure 6h**) because both low pH(Ghovanloo et al., 2018a; b; Peters et al., 2018) and CBD reduce current amplitudes to levels where differences in amplitudes could not be resolved above background noise.

**Figure 6.**
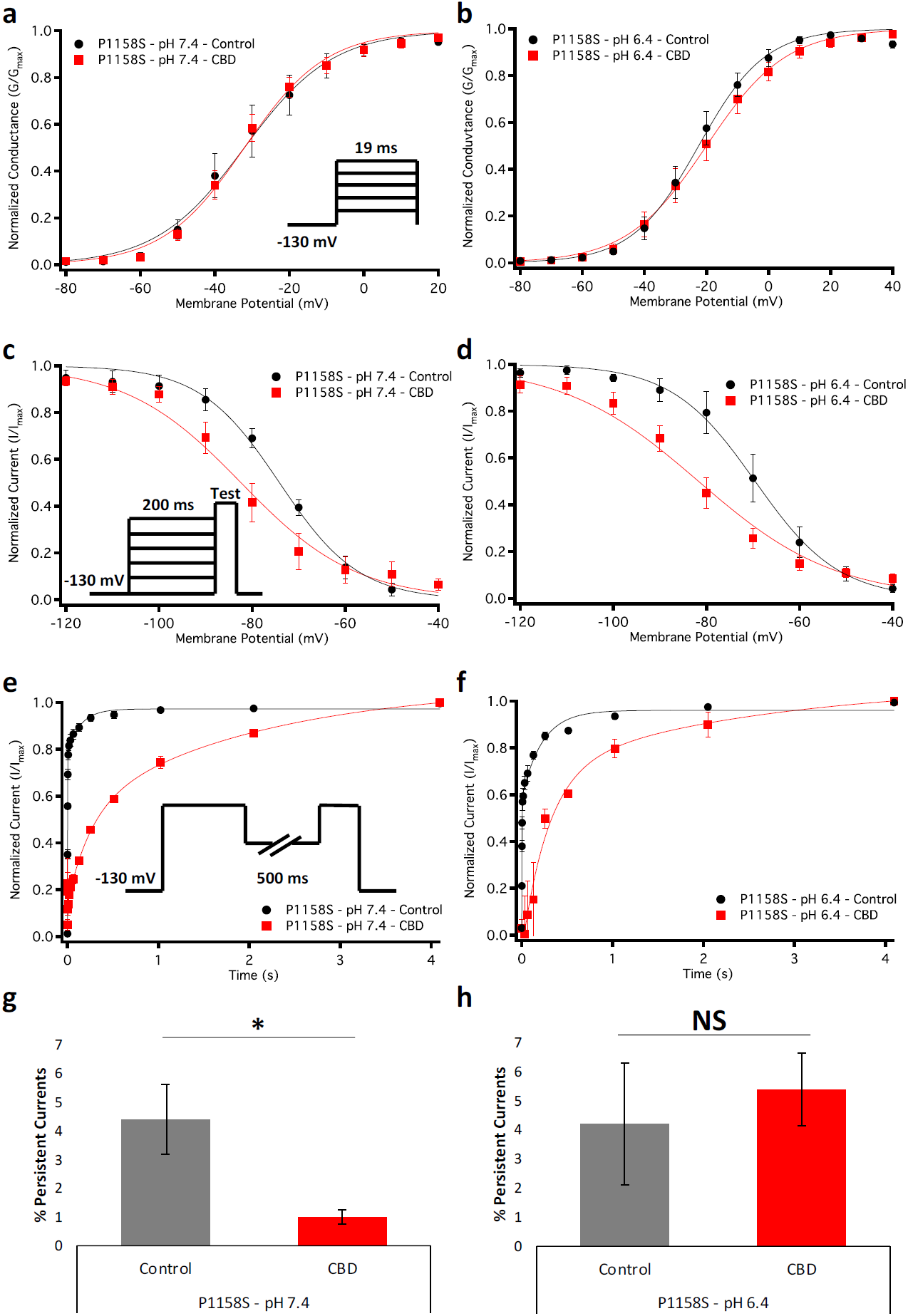
Effects of CBD (1 µM) on gating of a myotonia/hypoPP variant, P1158S. (**a-b**) Voltage-dependence of activation as normalized conductance plotted against membrane potential, at pH7.4 (Control: V_1/2_ = -30.0 ± 3.3 mV, z = 3.1 ± 0.2; CBD: V_1/2_ = -32.7 ± 3.6 mV, z = 2.9 ± 0.2; n = 7-8) and pH6.4 (Control: V_1/2_ = -23.0 ± 3.3 mV, z = 2.9 ± 0.2; CBD: V_1/2_ = -21.1 ± 3.3 mV, z = 2.5 ± 0.2; n = 8). (**c-d**) Voltage-dependence of SSFI plotted against membrane potential at pH7.4 (Control: V_1/2_ = -73.2 ± 2.6 mV, z = 2.9 ± 0.2; CBD: V_1/2_ = -83.0 ± 2.6 mV, z = 3.0 ± 0.3; n = 7) and pH6.4 (Control: V_1/2_ = -68.4 ± 3.0 mV, z = 2.7 ± 0.4; CBD: V_1/2_ = -81.7 ± 2.3 mV, z = 2.7 ± 0.3; n = 5-9). (**e-f**) Recovery from fast inactivation at 500 ms at pH7.3 (Control: *τ*_Fast_ = 0.0018 ± 0.006 s, *τ*_Slow_ = 0.15 ± 0.6 s; CBD: *τ*_Fast_ = 0.24 ± 0.07s; *τ*_Slow_ = 2.5 ± 0.6 s; n = 6-7) and pH6.4 (Control: *τ*_Fast_ = 0.065 ± 0.04 s, *τ*_Slow_ = 0.75 ± 0.4 s; CBD: *τ*_Fast_ = 0.13 ± 0.07 s; *τ*_Slow_= 0.62 ± 0.1 s; n = 4-7). (**g-h**) Persistent currents measured from a 200 ms depolarizing pulse to 0 mV from a holding potential of -130 mV at pH7.4 (Control: Percentage = 4.4 ± 1.2%; CBD: Percentage = 1.0 ± 0.2%; n = 4) and pH6.4 (Control: Percentage = 4.4 ± 2.1%; CBD: Percentage = 5.4 ± 1.2%; n = 5-6).

### AP model predicts that CBD reduces myotonia, but not hypoPP in the P1158S-pH assay

We used the gating changes from the patch-clamp experiments with WT and P1158S (both control and 1 µM CBD) to model the skeletal muscle AP(Cannon et al., 1993; Ghovanloo et al., 2018a). We ran the simulations using a 50 µA/cm^2^ stimulus. The simulation pulse started at 50 ms and stopped at 350 ms (**Figure 7**). During this pulse, the WT channels activated at 50 ms and fired a single AP. The channels remained inactivated until the stimulus was removed at 350 ms, and then the membrane potential recovered back to its resting value (**Figure 7a**). CBD reduced the AP amplitude (**Figure 7b**), consistent with CBD effects observed in different neuron types(Khan et al., 2018; Ghovanloo et al., 2018c). At pH6.4, P1158S displayed a continuous train of APs for the entire stimulation period. After the stimulus was removed, P1158S showed a progressive series of after-depolarizations of the membrane potential, characteristic of a myotonic burst (**Figure 7c**)(Cannon, 2015). Interestingly, the CBD-mediated shifts at pH6.4 in P1158S reduced the simulated AP amplitudes for the entirety of the pulse duration, delayed onset of first AP, consistent with CBD preventing Nav opening (as shown by measurements on peak Nav conductance in: (Ghovanloo et al., 2018c)), and abolished the post-pulse myotonic after-depolarizations (**Figure 7d**). At pH7.4, P1158S fired a single AP, followed by a period where membrane potential remained depolarized around −35 mV, even post-stimulus termination (**Figure 7e**). This inability to repolarize holds the Navs in an inactivated state, and is consistent with the periodic paralysis phenotype(Cannon, 2015). In contrast to the myotonic phenotype, CBD did not alleviate the hypoPP phenotype in our P1158S-pH *in-vitro/in-silico* assay (**Figure 7f**), consistent with its slowing of recovery from fast inactivation.

**Figure 7.**
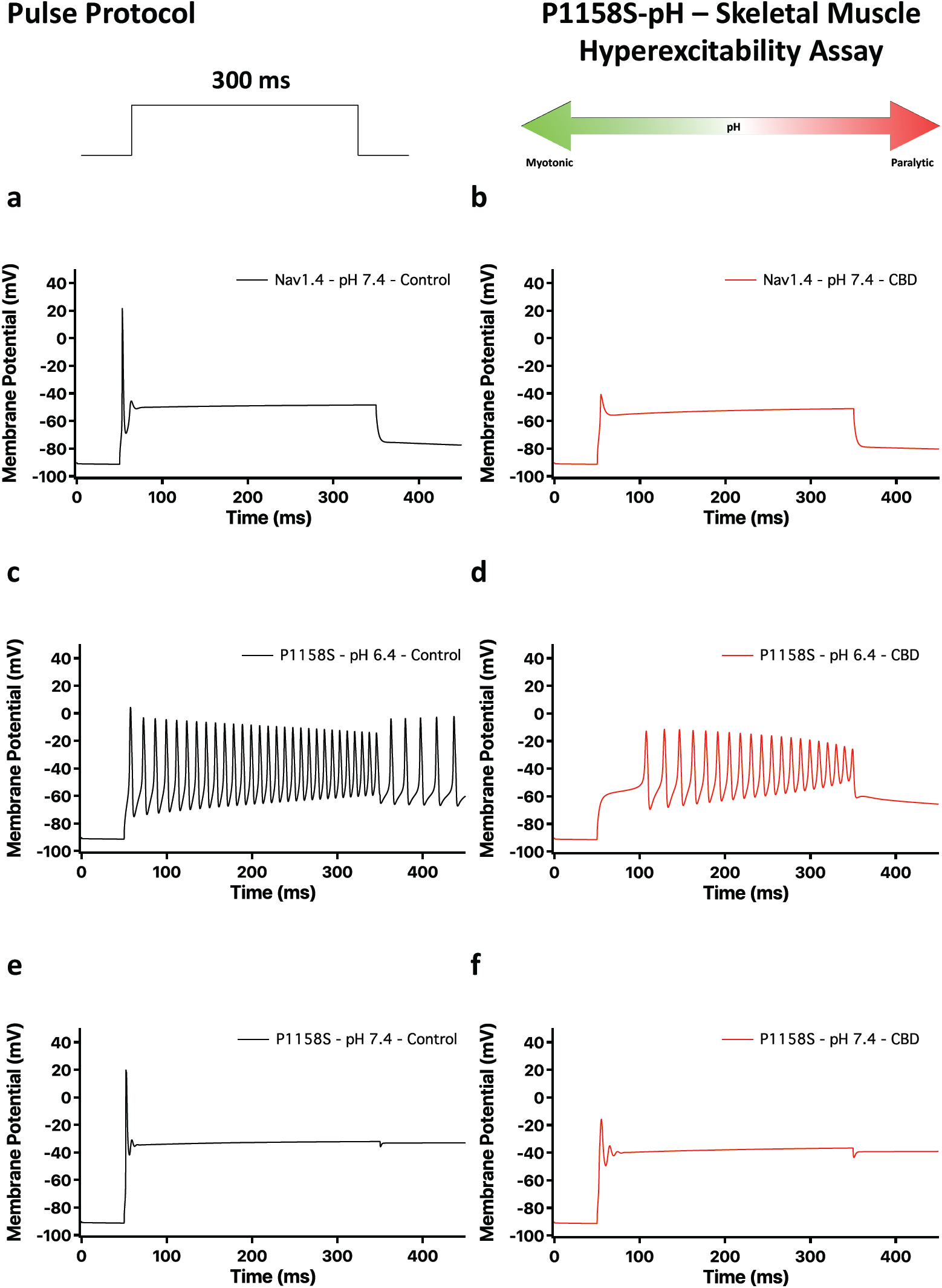
AP simulations of skeletal muscle action potentials in presence and absence of CBD, based on voltage-clamp data. Top of the figure show pulse protocol used for simulations, and a cartoon representation of P1158S-pH *in-vitro/in-silico* assay, where pH can be used to control the P1158S phenotype. (**a-b**) Show simulations in WT-Nav1.4 in presence and absence of CBD. (**c-d**) Show simulations of P1158S at pH6.4. (**e-f**) Show results from pH7.4.

### CBD reduces rat diaphragm muscle contraction at saturating concentrations

To survey and determine whether CBD reduces skeletal muscle contractions, we surgically removed rat diaphragm muscles and measured muscle contractions evoked by phrenic nerve stimulation. In **Figure 8a-b**, we show images of the diaphragm, cut into a hemi-diaphragm. We used electrodes to stimulate the phrenic nerve and measured the muscle contraction using a force transducer, at a saturating concentration of 100 µM of CBD, reasoning that if CBD reduces muscle contraction, a saturating concentration should provide a large enough response to detect any potential reduction in contraction. Our results suggested that CBD reduces the contraction amplitude to ∼60% of control (p<0.05) (**Figure 8c**). Next, we sought to determine whether a selective block of Nav channels also reduces skeletal muscle contraction using 300 nM tetrodotoxin (TTX), a saturating concentration of this potent blocker of selected Nav channels (IC_50_ ∼10-30 nM on TTX-sensitive channels(Hille, 2001)). TTX also reduced contraction to ∼20% of control (p<0.05) (**Figure 8c**). The remaining ∼20% contraction could be due to stimulation of voltage-gated calcium channels in transverse membranes that directly interact with ryanodine-sensitive calcium release in the SR that can initiate contraction(Catterall, 2011; TANABE et al., 1993; Catterall, 1991). Representative traces of muscle contraction in control, CBD, and TTX are shown in **Figure 8d-f**. These results show that a selective inhibition of Nav reduces skeletal muscle contraction, and that suggests that CBD’s reduction of muscular contraction could be due, at least in part, to its effect on Nav(Ghovanloo et al., 2018c). Therefore, our molecular *in-vitro* and *in-silico* data could have some physiological relevance.

**Figure 8.**
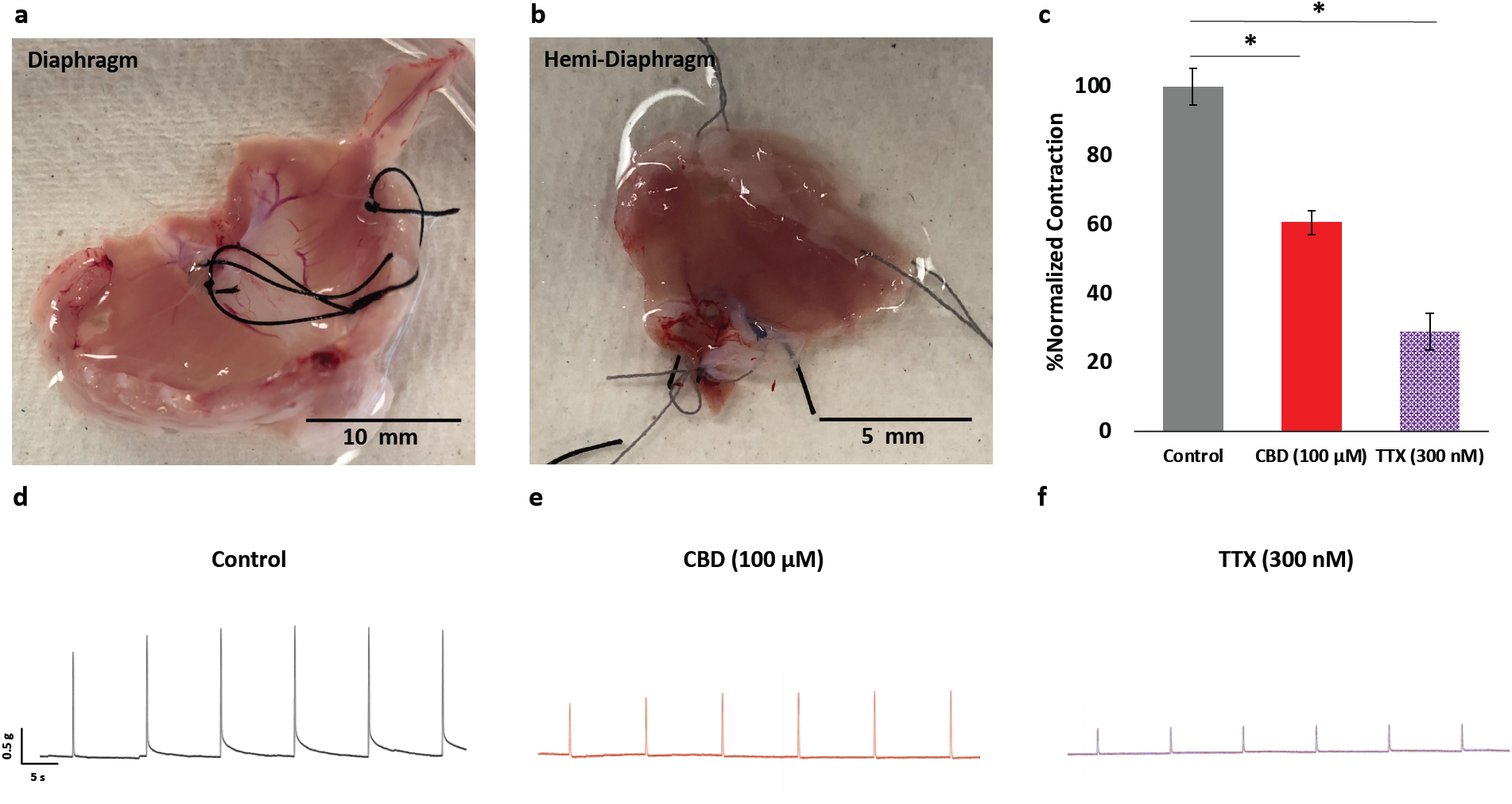
Effects of CBD on rat diaphragm contraction. (**a**)Image of dissected rat diaphragm muscle. (**b**) Image of rat diaphragm cut into a hemi-diaphragm, which was placed between electric plates that were used for electric stimulation. The subsequent muscle contractions were measured using a force transducer. (**c**) Normalized quantification of muscle contractions in CBD and TTX (Percentage of normalized contraction: Control = 100 ± 5.3%; CBD = 60.6 ± 3.5%; TTX = 28.9 ± 5.3%; n = 6-9). (**d-f**) Sample contraction traces across all three conditions.

## DISCUSSION

### Pathway and mechanism of Nav1.4 inhibition by CBD

Although CBD holds therapeutic promise(Devinsky et al., 2017; Ghovanloo et al., 2018c; Kaplan et al., 2017; Patel et al., 2016; Ross et al., 2008; Pumroy et al., 2019; Fouda et al., 2020) and has been approved for two seizure disorders, its mechanisms of action remain largely unknown. We previously described the effects of CBD on Nav, which are among its proposed targets(Ghovanloo et al., 2018c). CBD’s effects on neuronal Nav resemble the properties described for both amphiphilic compounds and traditional pore-blockers. That study provided foundational hypotheses about CBD’s mechanism of action on Nav. Here, we tested those ideas using a combination of *in-vitro, in-silico*, and *ex-vivo* techniques.

Amphiphiles, those molecules possessing both lipophilic and hydrophilic properties, often display non-selective modulatory effects on seemingly unrelated targets(Lundbæk et al., 2004; Lundbaek, 2005; Kapoor et al., 2019). The apparent diversity of targets is a by-product of amphiphiles modulating membrane elasticity(Lundbæk et al., 2004; Lundbaek, 2005; Kapoor et al., 2019). This modification is achieved by amphiphiles localizing at the solution–bilayer interface, which is made possible by having the compound’s polar group residing at the interface with the hydrophobic region, which then gets inserted into the bilayer core. This partitioning into the lipid bilayer alters membrane elasticity, and changes phase preference and curvature(Lundbæk et al., 2004; Lundbaek, 2005; Kapoor et al., 2019). The net effect of these alterations to the membrane for the bilayer-embedded Nav channel is a stabilized inactivated state(Lundbæk et al., 2004; Lundbaek, 2005; Kapoor et al., 2019).

We used MD simulations to ‘visualize’ CBD localization and its effects on the membrane. Interestingly, the MD prediction regarding localization, independently confirmed by NMR measurements of CBD in lipid vesicles, suggested CBD positioning between C8-C10. Our gramicidin-based functional assay suggested that CBD slightly changes membrane elasticity. This result is consistent with our previous findings, including CBD’s temperature-dependence (CBD effects were enhanced at lower temperatures), stabilized Nav inactivation, and non-selectivity of Nav inhibition(Ghovanloo et al., 2018c). Together, these findings suggest that CBD inhibition of Nav currents (and possibly other ionic currents) is, at least in part, mediated through changing lipid bilayer elasticity.

In this study, we further found that CBD had the opposite effect to Triton X-100 in GFA. Also, the magnitude change of quench rate was different between the two compounds at a given concentration; however, both compounds similarly hyperpolarized the Nav inactivation. These findings suggest that there could be at least two, maybe three mechanisms involved. The exact mechanisms through which CBD’s presence alter the lipid/Nav interactions should be further investigated in future studies.

The modulated receptor hypothesis suggests that resting-state block occurs when a compound enters from the lipid phase of the membrane into the LA binding site, whereas rapid open-state block happens when a compound enters the open pore from the cytosol(Hille, 1977; Hondeghem and Katzung, 1984). Pore-blockers can reach their binding site from the cytosolic side when the activation gate is open. A recent study showed that compounds can have direct access from the membrane phase to the LA site through channel fenestrations, culminating in resting-state block(Gamal El-Din et al., 2018).

We previously found that some characteristics of CBD inhibition of Nav are similar to classic pore-blockers(Ghovanloo et al., 2018c). Here, we tested CBD interactions inside the Nav1.4 LA site at rest. Destabilizing the LA site by the F1586A mutation, reduced CBD block of Nav1.4. This result is particularly notable since the LA site becomes a more favourable interaction site when the channel adopts a more inactivated state (a key reason for LAs’ strong state-dependence)(Ghovanloo and Ruben, 2020). Therefore, CBD’s reduced block in F1586A at rest could support the idea that CBD interacts with Nav at the pore. However, this does not indicate that the pore is the primary determinant of CBD inhibition, especially when CBD is compared to a traditional blocker like lidocaine, which is more affected by the F1586A mutation. However, it is also possible that CBD’s interaction is merely less dependent on F1586 than lidocaine, with the pore LA site being equally critical for both compounds.

Next, we reasoned that, if CBD blocks the pore, a likely path to reach the pore from the lipid phase would be through the Nav fenestrations (based on MD results, high LogP, and high lipid binding partitioning). We found that our fenestration-occluded Nav1.4-WWWW construct abolished resting-state block by CBD but not lidocaine or flecainide. This could be a consequence of the differences in hydrophobicity (and size/shape) between these three compounds. Because CBD is several orders of magnitude more hydrophobic than either lidocaine or flecainide, it may preferentially reach the pore through the fenestrations, whereas lidocaine and flecainide can reach the pore also from the cytoplasmic phase even if access through the fenestrations is blocked. This interpretation is consistent with the modulated receptor hypothesis (**Figure S5**).

Finally, we determined that, while CBD’s I_Na_ block occurs through its interactions inside the Nav1.4 pore, its stabilization of inactivation at least in part arises from modulating membrane elasticity. Both mechanisms contribute to CBD’s overall inhibition of Nav currents.

### Possible clinical applications for CBD in skeletal muscle disorders

Skeletal muscle hyperexcitability disorders have historically received less attention than disorders in other tissues, including the brain. Drugs most commonly used for myotonia include compounds developed for other conditions, such as anti-convulsants and anti-arrhythmics(Alfonsi et al., 2007; Trip et al., 2008), which may cause unwanted, off-target side-effects. Hence, another therapeutic approach has been lifestyle modifications. For instance, myotonic patients may modify their lifestyles to avoid triggers like potassium ingestion or cold temperatures. Treatment of hypoPP is usually achieved using oral potassium ingestion and by avoiding dietary carbohydrates and sodium. During hypokalemia, increasing K^+^ levels may reduce membrane depolarization and shift the resting potential to more negative potentials. Acetazolamide or dichlorphenamide may be useful; however, these compounds can exacerbate symptoms(Torres et al., 1981; Tawil et al., 2000; Sternberg et al., 2001; Venance et al., 2004). There is a need for new treatments for these conditions.

Cannabinoids have long been used to alleviate muscular problems(Borgelt et al., 2013; Baker et al., 2000). In this study, we show CBD reduces skeletal contraction in rat diaphragm muscle. As CBD is a poly-pharmacology compound, we cannot state with certainty that the observed contraction reduction is due to I_Na_ inhibition alone, but as demonstrated with the TTX results, I_Na_ block is sufficient to reduce contraction, meaning that CBD’s activity at Nav1.4 could be a part of the mechanism in this reduction. Another caveat is that we cannot exclude the possibility of a phrenic nerve independent (i.e. direct muscle) stimulation resulting in muscle contraction in our myography experiments. The overall mechanism suggested by our results is summarized in **Figure 9**.

**Figure 9.**
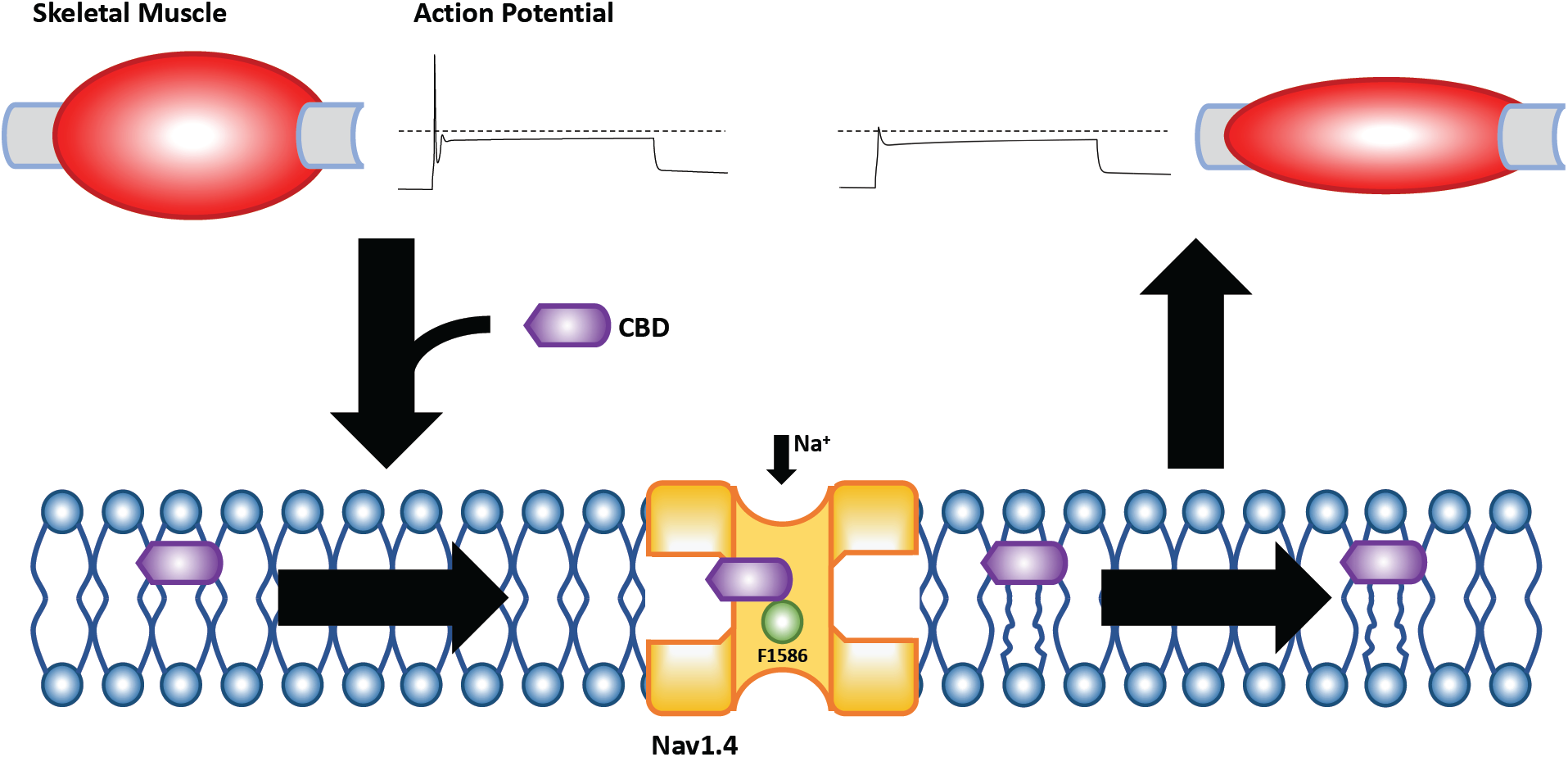
Pathway of skeletal muscle inhibition via Nav1.4. This is a cartoon representation of the mechanism and pathway through which CBD inhibits Nav1.4. Once CBD is exposed to the skeletal muscle, given its high lipophilicity, the majority of it gets inside the sarcolemma. Upon entering the sarcolemma, it localizes in the middle regions of the leaflet, and travels through the Nav1.4 fenestrations into the pore. Inside the pore mutation of the LA F1586A reduces CBD inhibition. CBD also alters the membrane elasticity, which promotes the inactivated state of the Nav channel, which adds to the overall CBD inhibitory effects. The net result is a reduced electrical excitability of the skeletal muscle, which - at least in part - contributes to a reduction in muscle contraction.

To explore a possible use for CBD in myotonia and hypoPP, we tested it in an *in-vitro/in-silico* assay. Our results suggest that CBD may alleviate the myotonic but not the hypoPP phenotype. One caveat is that these predictions are based in part on computer simulations. However, from a theoretical perspective, most Nav1.4 mutations that cause myotonia do so by changing conventional channel gating (e.g. activation, inactivation, persistent currents); hypoPP mutants are due to pathogenic gating pore currents associated with the VSD, so it is conceivable for a compound like CBD to alleviate myotonic behavior, but not hypoPP.

In conclusion, our results suggest that CBD inhibition of Nav has at least two components: altered membrane elasticity and pore block. Nav1.4 inhibition could contribute to CBD reducing skeletal muscle contractions and may have potential therapeutic value against myotonia (**Figure 9**). From a broader perspective, our proposed mechanism may hold true for other compounds that are similar to CBD in modulating Navs or other channels with similar structures.

## MATERIALS AND METHODS

### Rat diaphragm preparation

Four 4-week old male Sprague Dawley rats (Charles River, Raleigh site) were euthanized. The rat phrenic hemi-diaphragm preparation was isolated according to the method described by Bulbring (1946)(BULBRING, 1946). A fan-shaped muscle with an intact phrenic nerve was isolated from the left side and transferred to a container with Krebs’ solution (NaCl 95.5, KCl 4.69, CaCl_2_ 2.6, MgSO_4_.7H_2_O 1.18, KH_2_PO_4_ 2.2, NaHCO_3_ 24.9, and glucose 10.6 mM) and aerated with carbogen (95% oxygen and 5% carbon dioxide). All experimental protocols were approved by the Animal Care and Use Committees. Contraction experiment was performed using a Radnoti Myograph system.

### Molecular docking

Docking of CBD into the cryo-EM structure of hNav1.4 (PDB ID: 6AGF was carried out using Autodock Vina(Trott and Olson, 2010). CBD was downloaded in PDB format from Drugbank(Wishart et al., 2018). To dock CBD into Nav1.4 a large search volume of 32 Å x 44 Å x 26 Å was considered, that enclosed nearly the whole of the pore domain and parts of VSD. This yielded the top 9 best binding poses of CBD ranked by mean energy score.

### MD simulation systems preparation

We ran two different sets of MD simulations, a first consisting of CBD interacting with model POPC membranes, and a second consisting of CBD interacting with the hNav1.4 channel embedded in its POPC/solution environment.

First, a homogenous lipid bilayer consisting of 188 POPC molecules was prepared using the CHARMM-GUI Membrane builder(Jo et al., 2008; Lee et al., 2016; Wu et al., 2014). Three different systems were created: one with two CBD molecules, each one placed in each leaflet of the bilayer (System 1), one with three CBDs, all of them placed in the upper leaflet (System 2) and one with six CBDs, of which three were placed in the upper leaflet and three in the lower leaflet (System 3). CBD was placed manually into the bilayer, with the polar headgroup of CBD facing the lipid headgroups. Lipid molecules with at least one atom within 2 Å of a CBD were manually deleted. A control simulation without any CBD was also prepared (System 0). The system was hydrated by adding two ∼25 Å layers of water to both sides of the membrane. Lastly, 150 mM NaCl was added (30 Na+ and 30 Cl^−^). The simulation systems are summarized in **Table 1**. This system was defined as the lipid-CBD

Second, hNav1.4 and the best docked position of CBD obtained from Autodock Vina was used as a starting structure. The starting system was embedded into POPC lipid bilayer. The system was hydrated by adding two ∼25 Å layers of water to both sides of the membrane. Lastly, the system was ionized with 150 mM NaCl. This system is defined as the Nav1.4-CBD-lipid system.

### MD simulations

The CHARMM36 forcefield was used to describe the protein, lipid bilayer, and the ions(Klauda et al., 2010). CBD was parameterised using the SWISS-PARAM software(Zoete et al., 2011). The TIP3P water model was used to describe the water molecules. The systems were minimised for 5000 steps using steepest descent and equilibrated with constant number of particles, pressure and temperature (NPT) for at least 450 ps for the lipid-CBD system and 36 ns for the Nav1.4-CBD-lipid system, during which the position restraints were gradually released according to the default CHARMM-GUI protocol. During equilibration and production, a time step of 2 fs was used, pressure was maintained at 1 bar through Berendsen pressure coupling, temperature was maintained at 300 K through Berendsen temperature coupling with the protein, membrane and solvent coupled and LINCs algorithm(Hess et al., 1997) was used to constrain the bonds containing hydrogen. For long range interactions, periodic boundary conditions and particle mesh Ewald (PME) were used. For short range interactions, a cut-off of 12 Å was used. Finally, unrestrained production simulations were run for 150 ns for each of the lipid-CBD system and 10 ns for the Nav1.4-CBD-lipid system, using Parinello-Rahaman pressure coupling(Parrinello and Rahman, 1981) and Nose-Hoover temperature coupling(Nosé, 1984). Simulations were performed using GROMACS 2018.4(Abraham et al., 2015)

### ABMD simulations

Adiabatic biased molecular dynamics (ABMD)(Marchi and Ballone, 1999) simulations were performed using GROMACS 2018.4 (Abraham et al., 2015) patched with Plumed-2.5.1(Tribello et al., 2014) to study the entrance pathway of CBD into its docking site in hNav1.4. ABMD is a simulation method in which a time dependent biasing harmonic potential is applied to drive the system towards a target system. along a predefined collective variable. Whenever the system moves closer towards the target system along the collective variable, the harmonic potential is moved to this new position, resulting in pushing the system towards the final state. The bias potential was applied along the distance between the center of masses of CBD and F1586. Two types of biasing potentials were considered: one along the y-component of distance and other along all components of distance.

### ^2^H NMR lipid analysis

1-palmitoyl-2-oleoyl-sn-glycero-3-phosphocholine (POPC-d31, sn-1 chain perdeuterated) was obtained from Avanti Polar Lipids (Alabaster, AL). The POPC-d31:CBD sample was prepared with ∼50 mg lipid and 3.4 mg of CBD for a ratio of POPC/CBD 8:2. The two samples, pure POPC-d31 and POPC-d31:CBD (8:2), were dissolved in Bz/MeOH 4:1 (v/v) and freeze-dried. After hydration with excess amounts of deuterium-depleted water (ddw), five freeze-thaw-vortex cycles were done between liquid nitrogen (−196°C) and 60°C to create multilamellar dispersions (MLDs).

Deuterium ^2^H NMR experiments were performed on a TacMag Scout spectrometer at 46.8 MHz using the quadrupolar echo technique1. The spectra were produced from ∼20,000 two-pulse sequences. 90° pulse lengths were set to 3.1 μs, inter-pulse spacing was 50 μs, dwell time was 2 μs, and acquisition delays were 300 ms. Data were collected using quadrature with Cyclops eight-cycle phase cycling. The spectra were dePaked to extract the smoothed order parameter profiles of the POPC sn-1 chain in the presence or absence of CBD. Samples were run at 20, 30, and 40°C, left to equilibrate at each temperature for 20 mins before measurements were taken.

### Gramicidin-fluorescence membrane elasticity assay

1,2-dierucoyl-sn-glycero-3-phosphocholine (DC_22:1_PC) were from Avanti Polar Lipids (Alabaster, AL). CBD was from Sigma-Aldrich (St. Louis, MO). 8-Aminonaphthalene-1,3,6-trisulfonate (ANTS) was from Invitrogen Life Technologies (Grand Island, NY). Gramicidin D was from (Sigma Aldrich).

GFA: Large unilamellar vesicles (LUVs) were made from DC_22:1_PC as described previously(Rusinova et al., 2015). Briefly, phospholipids in chloroform and gramicidin (gA) in methanol (1000:1 lipid:gA weight ratio) were mixed. Quench rates were obtained by fitting the quench time course from each mixing reaction with a stretched exponential(Ingólfsson et al., 2010):

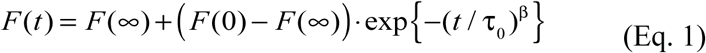

and evaluating the quench rate at 2 ms (the instrumental dead time is ∼1.5 ms):

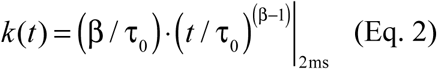

To test drug effects on the lipid bilayer CBD was equilibrated with the LUVs for 10 min at 25 °C before acquiring quench time courses. Each measurement consisted of (4 – 8) individual mixing reactions, and the rates for each mixing reaction were averaged and normalized to the control rate in the absence of drug.

### Cell culture

Chinese Hamster Ovary (CHOK1) cells were transiently co-transfected with cDNA encoding eGFP and the β1-subunit and either WT-Nav1.4 (GenBank accession number: NM_000334) or any of our mutant *α*-subunits. Transfection was done according to the PolyFect transfection protocol. After each set of transfections, a minimum of 8-hour incubation was allowed before plating on sterile coverslips.

### Patch-clamp

Whole-cell patch-clamp recordings were performed in an extracellular solution containing (in mM): 140 NaCl, 4 KCl, 2 CaCl_2_, 1 MgCl_2_, 10 HEPES or MES (pH6.4). Solutions were adjusted to pH6.4 and 7.4 with CsOH. Pipettes were filled with intracellular solution, containing (in mM): 120 CsF, 20 CsCl, 10 NaCl, 10 HEPES. In some experiments lower sodium concentration of 1 mM (intracellular) was used to boost driving force, and hence current size. All recordings were made using an EPC-9 patch-clamp amplifier (HEKA Elektronik, Lambrecht, Germany) digitized at 20 kHz via an ITC-16 interface (Instrutech, Great Neck, NY, USA). Voltage-clamping and data acquisition were controlled using PatchMaster/FitMaster software (HEKA Elektronik, Lambrecht, Germany) running on an Apple iMac. Current was low-pass-filtered at 10 kHz. Leak subtraction was performed automatically by software using a P/4 procedure following the test pulse. Giga-ohm seals were allowed to stabilize in the on-cell configuration for 1 min prior to establishing the whole-cell configuration. Series resistance was less than 5 MΩ for all recordings. Series resistance compensation up to 80% was used when necessary. All data were acquired at least 1 min after attaining the whole-cell configuration. Before each protocol, the membrane potential was hyperpolarized to −130 mV to ensure complete removal of both fast inactivation and slow-inactivation. All experiments were conducted at 22 ± 2 °C. Analysis and graphing were done using FitMaster software (HEKA Elektronik) and Igor Pro (Wavemetrics, Lake Oswego, OR, USA). All data acquisition and analysis programs were run on an Apple iMac (Apple Computer).

Some cDNA constructs produced small ionic currents. To ensure, the recorded currents were indeed construct-produced currents and not endogenous background currents, untransfected cells were patched and compared to transfected cells. The untransfected CHOK1 cells, which were exclusively used for cDNA expression, produced no endogenous sodium currents.

### Activation protocol

To determine the voltage-dependence of activation, we measured the peak current amplitude at test pulse potentials ranging from −100 mV to +80 mV in increments of +10 mV for 20 ms. Channel conductance (G) was calculated from peak I_Na_:

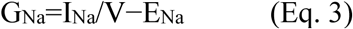

where G_Na_ is conductance, I_Na_ is peak sodium current in response to the command potential V, and E_Na_ is the Nernst equilibrium potential. Calculated values for conductance were fit with the Boltzmann equation:

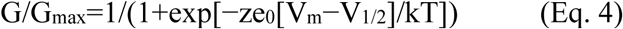

where G/G_max_ is normalized conductance amplitude, V_m_ is the command potential, z is the apparent valence, e_0_ is the elementary charge, V_1/2_ is the midpoint voltage, k is the Boltzmann constant, and T is temperature in K.

### Steady-state fast inactivation protocol

The voltage-dependence of fast inactivation was measured by preconditioning the channels to a hyperpolarizing potential of −130 mV and then eliciting pre-pulse potentials that ranged from −170 to +10 mV in increments of 10 mV for 500 ms, followed by a 10 ms test pulse during which the voltage was stepped to 0 mV. Normalized current amplitudes from the test pulse were fit as a function of voltage using the Boltzmann equation:

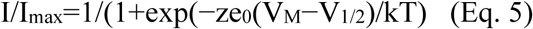

where *I*_max_ is the maximum test pulse current amplitude.

### Persistent current protocol

Persistent current was measured between 145 and 150 ms during a 200 ms depolarizing pulse to 0 mV from a holding potential of -130 mV. Pulses were averaged to increase signal-to-noise ratio.

### Recovery from fast inactivation protocol

Channels were fast inactivated during a 500 ms depolarizing step to 0 mV, and recovery was measured during a 19 ms test pulse to 0 mV following a −130 mV recovery pulse for durations between 0 and 4 s. Time constants of fast inactivation recovery showed two components and were fit using a double exponential equation:

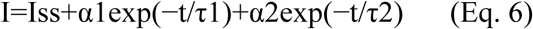

where *I* is current amplitude, *I*_ss_ is the plateau amplitude, α_1_ and α_2_ are the amplitudes at time 0 for time constants τ_1_ and τ_2_, and *t* is time.

### Isothermal titration calorimetry

The peptide with the following sequence: SYIIISFLIVVNM (from Nav1.4 DIV-S6, residues 1580-1592) was synthesized by GenScript. It was solubilized in DMSO and diluted to a final concentration of 1 mM with the final buffer containing by percentage: 10% DMSO, 60% acetonitrile, 30% ITC buffer. Acetonitrile was required to solubilize the peptide. The ITC buffer contained 50 mM HEPES pH7.2 and 150 mM KCl. CBD and Lidocaine were each solubilized in DMSO and diluted to a final concentration of 40 mM and 100 mM, respectively in the same final buffer as the peptide. Each titrant was injected into the peptide containing sample cell 13 times each with a volume of 3 µM with the exception of the first injection which was 0.4 µM. Stirring speed was set at 750 rpm.

### Action potential modeling

Skeletal AP modeling was based on a model developed by Cannon et al., (1993). All APs were programmed and run using Python. The modified parameters were based on electrophysiological results obtained from whole-cell patch-clamp experiments(Cannon et al., 1993). The model accounted for activation voltage-dependence, SSFI voltage-dependence, and persistent I_Na_. The WT pH7.4 model uses the original parameters from the model. P1158S models were programmed by shifting parameters from the original Cannon model by the difference between the values in P1158S experiments at a given pH/CBD(Ghovanloo et al., 2018a; Cannon et al., 1993).

### Statistics

A one-factor analysis of variance (ANOVA) or t-test were, when appropriate, were used to compare the mean responses. *Post-hoc* tests using the Tukey Kramer adjustment compared the mean responses between channel variants across conditions. A level of significance α=0.05 was used in all overall *post-hoc* tests, and effects with p-values less than 0.05 were considered to be statistically significant. All values are reported as means ± standard error of means (SEM) for *n* recordings/samples. Power analysis with α=0.05 was performed to yield sufficient *n* size for each experiment. Analysis was performed in JMP version 14.

## Funding and acknowledgements

This work was supported by grants from Natural Science and Engineering Research Council of Canada and the Rare Disease Foundation to PCR and M-RG (CGS-D: 535333-2019 & MSFSS: 546467-2019), a MITACS Accelerate fellowship in partnership with Xenon Pharma, Inc to M-RG (IT10714), and a MITACS Elevate fellowship in partnership with Akseera Pharma, Inc. to MAF, grants from SciLifeLab and the Swedish Research Council to LD (VR 2018-04905), and a grant from the US National Institutes of Health to OSA (GM021342). The MD simulations were performed on resources provided by the Swedish National Infrastructure for Computing (SNIC) at PDC Centre for High Performance Computing (PDC-HPC). We thank Drs. Colin Peters, Eric Lin, and Nina Weishaupt for their contributions.

## Authors contributions

M-RG assembled data, performed patch-clamp experiments, assisted in ITC experiments, action potential modeling/simulations, functional assay development, data analysis, figure making, wrote manuscript, data interpretation, and assisted to conceiving of experiments. KC and TSB performed MD simulations and docking. MAF performed myography. KR performed ITC, assisted in mutagenesis, and various experimental conceptualizations. RR performed GFA. TP performed NMR. KN performed diaphragm preparation. ARW helped with NMR. DP assisted with myography. JT, OSA, LD, SJG, and PCR conceived the experiments and revised the manuscript critically. All co-authors edited the manuscript.

## SUPPORTING INFORMATION SI FIGURES

**Figure S1.**
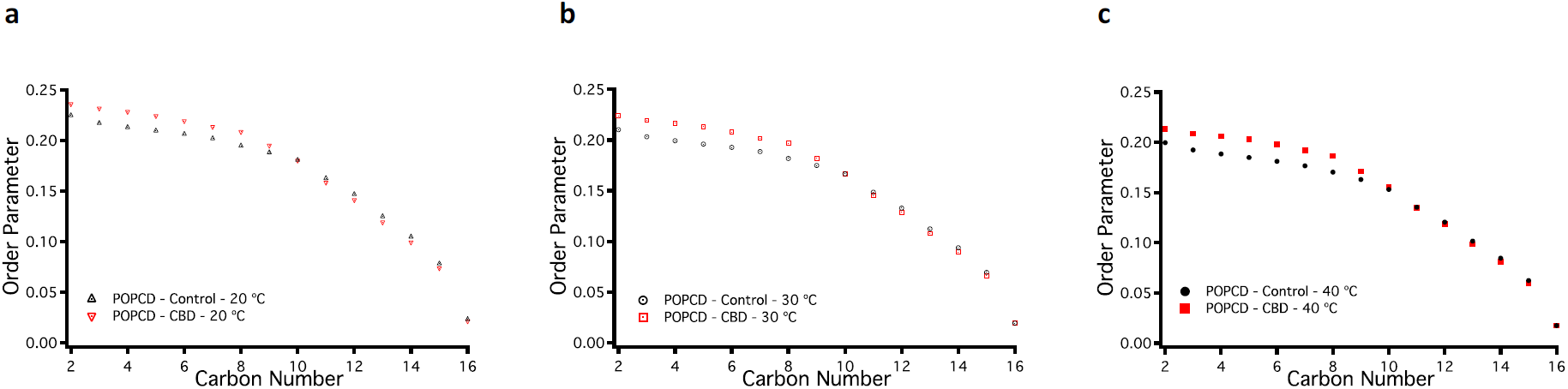
^2^H NMR at different temperatures. (**a-c**) Order parameters associated with POPC membranes at 20, 30, and 40 °C.

**Figure S2.**
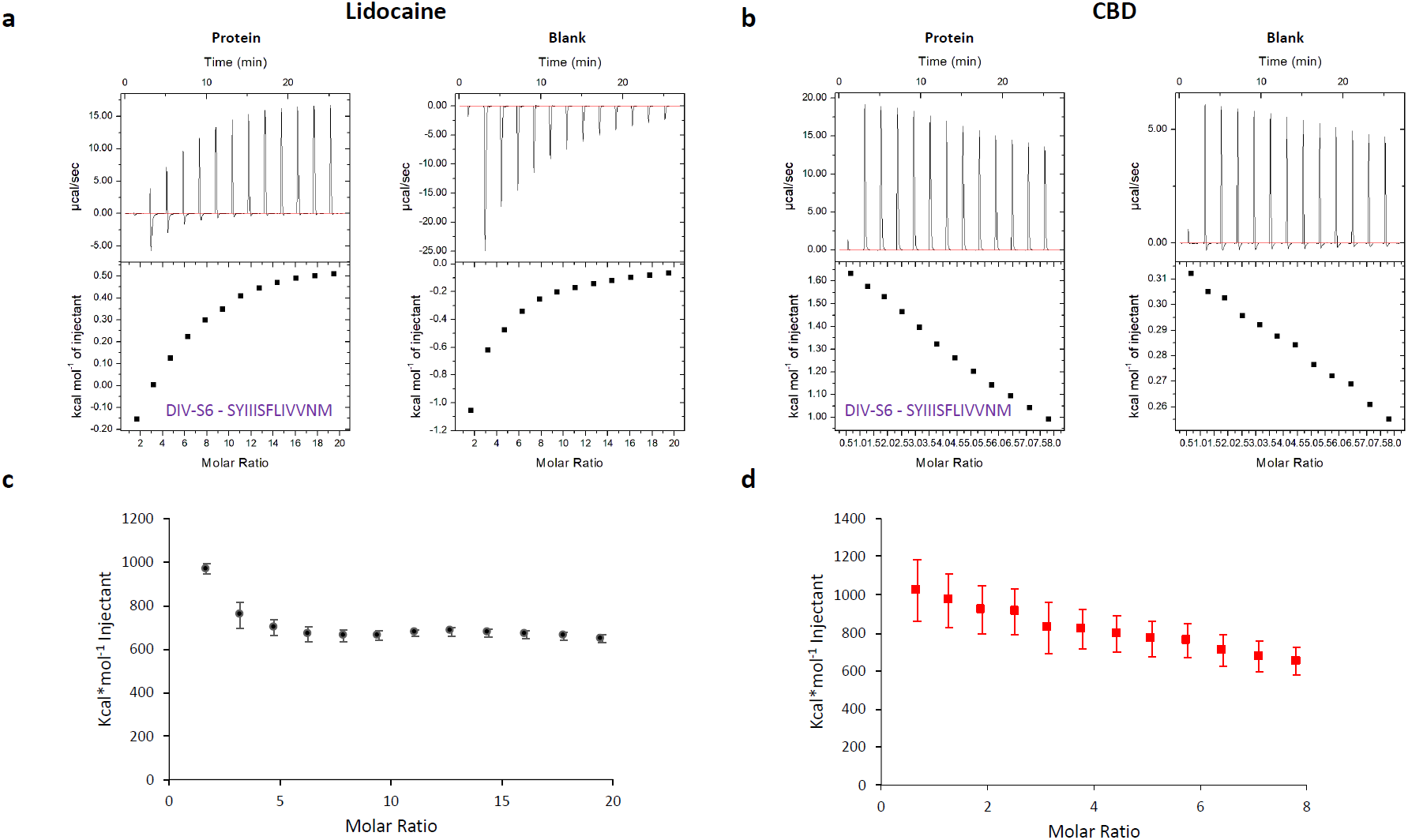
CBD interactions with DIV-S6, using isothermal titration calorimetry (ITC). (**a**) Representative ITC traces for titration of 100 mM lidocaine into 1 mM peptide or blank buffer. The heat signal, once the binding is saturated is the same as the blank if the blank was only measuring the interaction if lidocaine with the solution. The blank measures 3 interactions, interactions between solute molecules, solute and lidocaine and lidocaine and lidocaine. As more lidocaine is added with each injection the solution is changed. This makes the heat of interaction with in-blank trace different from beginning to end. This change is due to a change in amount of lidocaine in the solution. Because this change is progressive with each injection and that the injections into the peptide are the same volume, subtraction was used as a means to quantify lidocaine peptide interaction. (**b**) Representative ITC traces for titration of 40 mM CBD into 1 mM peptide or blank buffer. (**c**) The blank condition subtracted heat of titration in protein condition is shown for lidocaine, and (**d**) CBD. A peak heat of 968.0 ± 23.4 kcal*mol^-1^ was seen for lidocaine titration and a peak heat of 1022.2 ± 160.6 kcal*mol^-1^ was seen for the CBD titration (n = 3-4).

**Figure S3.**
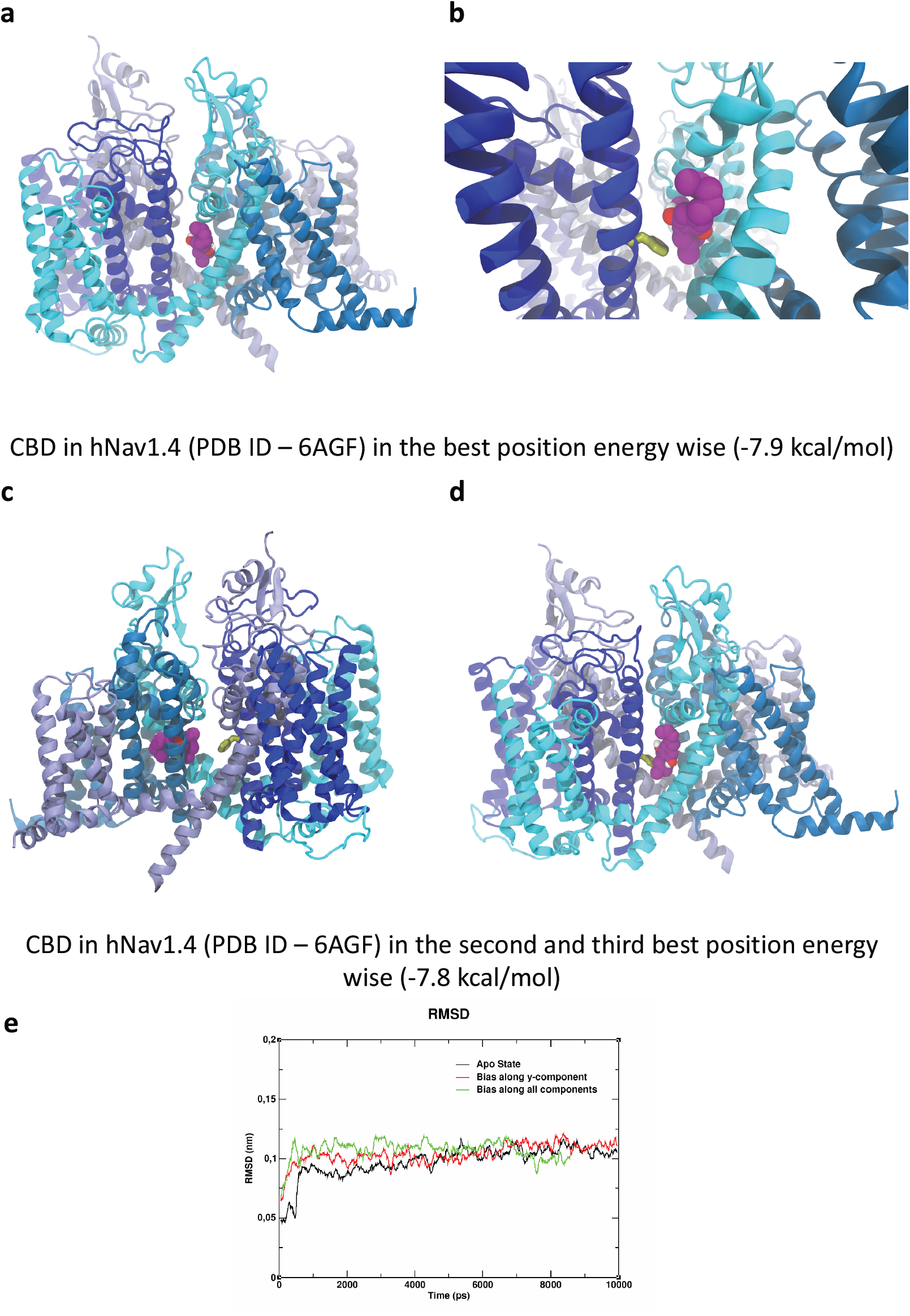
Nav1.4 fenestration interactions with CBD. (**a-d**) Shows CBD posed in the human Nav1.4 structure using molecular docking. (**e**) RMSD of the fenestration residues as a function of time in the absence (black) and the presence of CBD passing through the fenestration (red and green, two different simulation parameter sets). The similar RMSD profiles show that CBD’s passage does not distort the structural integrity of the fenestration.

**Figure S4.**
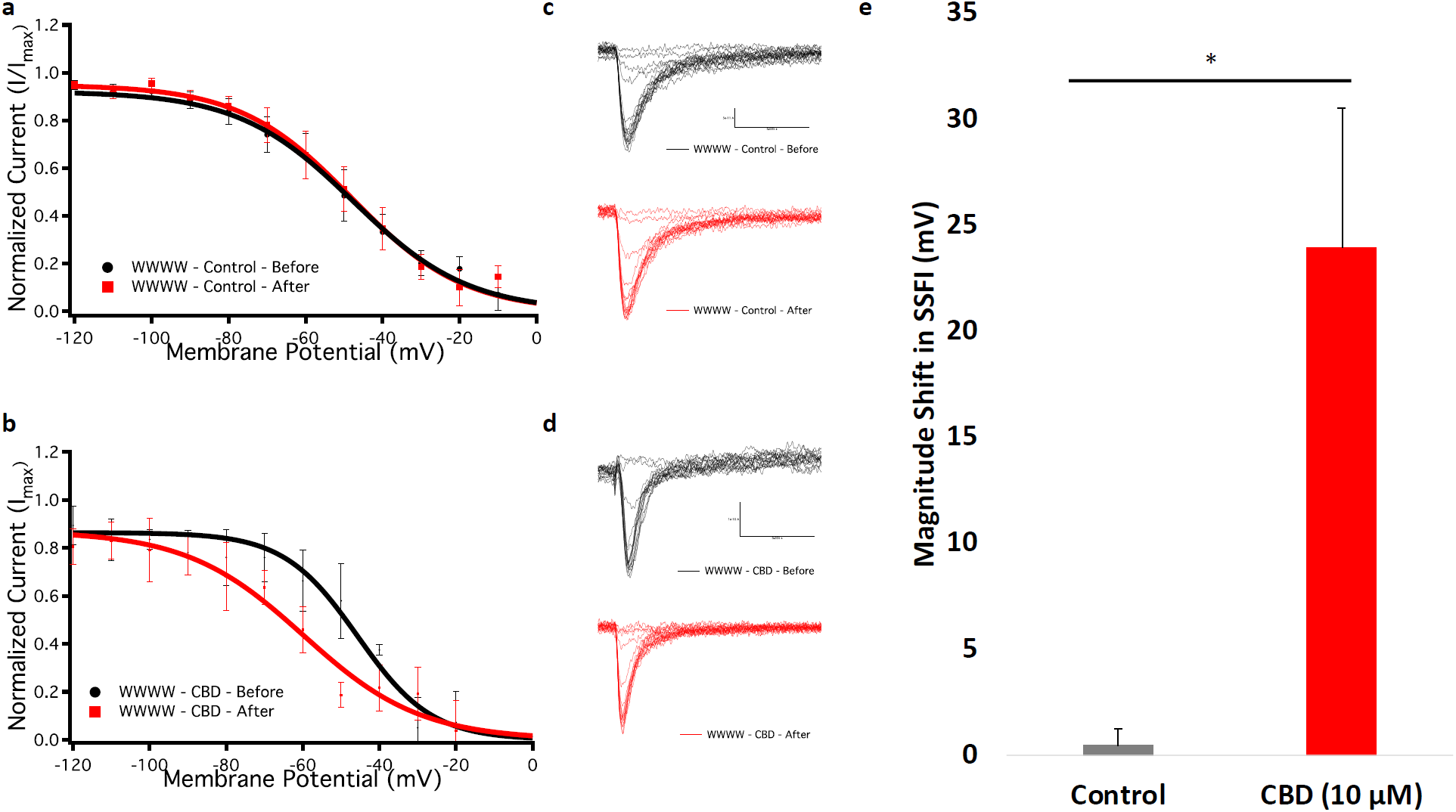
CBD stabilizes inactivation in the fenestration-occluded construct. (**a-b**) Show voltage-dependence of SSFI before and after (**a**) control (extracellular (ECS) solution) and CBD (10 µM) in WWWW construct (Before control: V1/2 = -54.7 ± 5.1 mV, After control: V_1/2_ = -54.2 ± 5.4 mV, Before CBD: V_1/2_ = -48.8 ± 8.8 mV, After CBD: V_1/2_ = -72.7 ± 5.7 mV, n = 3-6). The ECS experiment was performed to ensure that hyperpolarization shifts in the CBD condition are not due to possible confounding effects associated with fluoride in the internal (CsF) solutions. (**c-d**) Show representative families of inactivating currents before and after perfusion. CBD does not block peak currents but shifts the SSFI curve to the left. (**e**) Show averaged shift in the midpoint of SSFI before and after perfusion.

**Figure S5.**
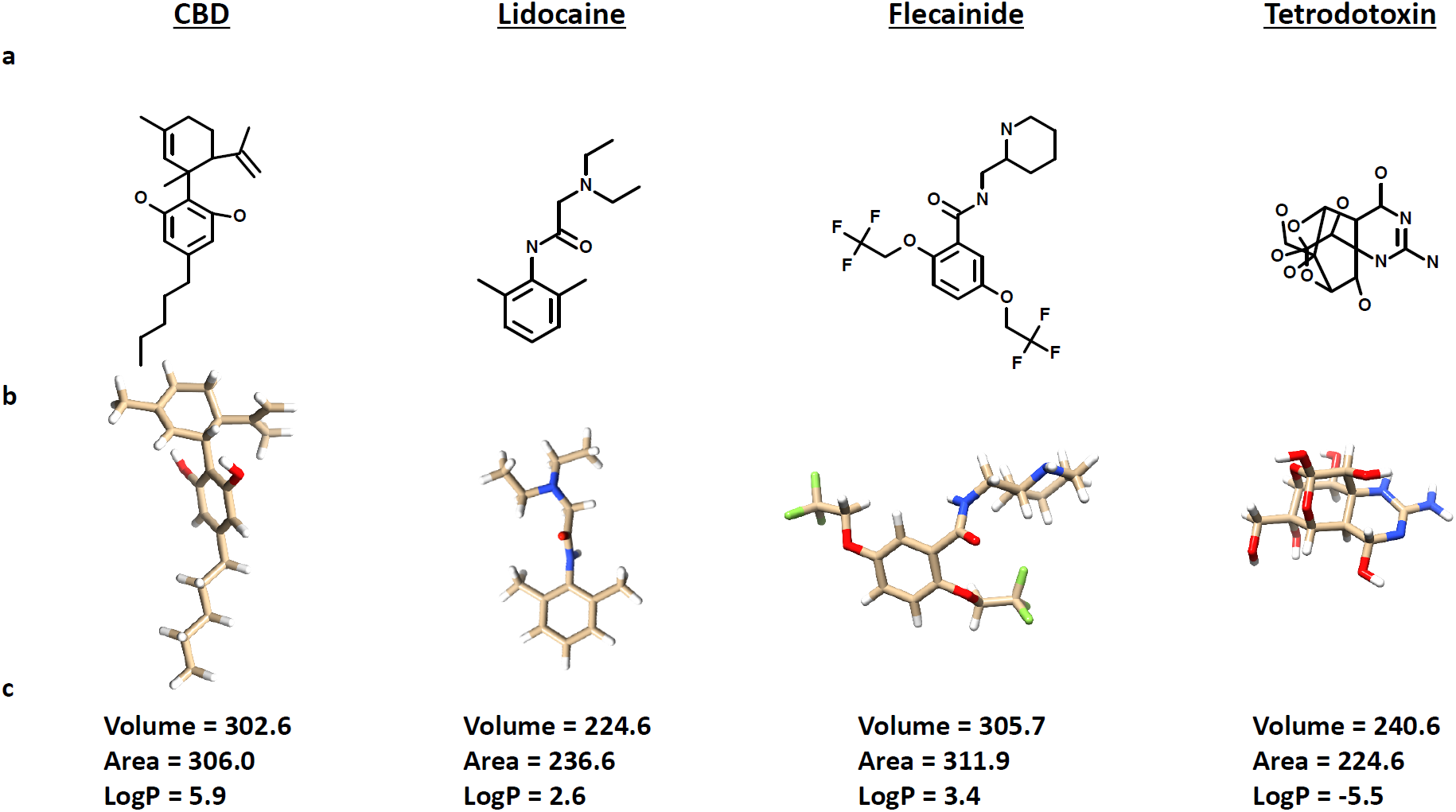
Comparison between some of the relevant physicochemical properties of the compounds used in this study. (**a**) Chemical structures of the compounds used in this study. (**b**) Three dimensional structures of the compounds. (**c**) Volume (Å^3^) and area (Å^2^) for each compound was calculated using UCSF Chimera. LogP values are obtained from ChEMBL database. CBD, lidocaine, and flecainide all interact with the LA site inside the Nav pore. TTX interacts with the outer selectivity filter of the Nav pore. CBD is several times more hydrophobic than the other compounds. CBD is larger than lidocaine and slightly smaller than flecainide.

## SI MOVIES

**Movie S1-**CBD localization inside POPC membrane.

**Movie S2-**ABMD simulation show that CBD passes through the fenestration of Nav1.4 (bias applied along y-component of distance).

**Movie S3-**ABMD simulation show that CBD passes through the fenestration of Nav1.4 (bias applied along all components of distance).

